# Bio-friendly long-term subcellular dynamic recording by self-supervised image enhancement microscopy

**DOI:** 10.1101/2022.11.02.514874

**Authors:** Guoxun Zhang, Xiaopeng Li, Yuanlong Zhang, Xiaofei Han, Xinyang Li, Jinqiang Yu, Boqi Liu, Jiamin Wu, Li Yu, Qionghai Dai

**Author notes:** These authors contributed equally to this work (Guoxun Zhang, Xiaopeng Li, Yuanlong Zhang). Correspondence (J.W.), (L.Y.), (Q.D.).

## Abstract

Fluorescence microscopy has become an indispensable tool for revealing the dynamic regulations of cells and organelles in high resolution noninvasively. However, stochastic noise inherently restricts the upper bonds of optical interrogation quality and exacerbates the observation fidelity in encountering joint demand of high frame rate, long-term, and low photobleaching and phototoxicity. Here, we propose DeepSeMi, a self-supervised-learning-based denoising framework capable of increasing SNR by over 12 dB across various conditions. With the introduction of newly designed eccentric blind-spot convolution filters, DeepSeMi accomplished efficacious denoising requiring no clean data as references and no compromise of spatiotemporal resolution on diverse imaging systems. The computationally 15-fold multiplied photon budget in a standard confocal microscope by DeepSeMi allows for recording organelle interactions in four colors and high-frame-rate across tens of thousands of frames, monitoring migrasomes and retractosomes over a half day, and imaging ultra-phototoxicity-sensitive *Dictyostelium* cells over thousands of frames, all faithfully and sample-friendly. Through comprehensive validations across various cells and species over various instruments, we prove DeepSeMi is a versatile tool for reliably and bio-friendly breaking the shot-noise limit, facilitating automated analysis of massive data about cell migrations and organelle interactions.

## Introduction

The magnificence of the harmonically orchestrated systems, organs, tissues, and cells attracts people to exploit the mystery of life[1, 2]. Among the various phenotypic activities and processes, organelles interact with one another and the cytoskeleton to synergistically execute various physiological functions that support the functioning of living beings. Such gorgeous patterns reflect live organelles of complex, dynamic, and interplay in highly dynamic yet organized interactions capable of orchestrating complex cellular functions[3]. Thereby, visualizing the functionality and complexity of organelles in their native states requires high spatiotemporal resolution observation without perturbing these physiologically presented regulations in a long term.

Standing in the center of approaches dedicated to probing and deciphering the micro world is the non-invasive fluorescent microscope capable of high spatiotemporal resolution[4] and good protein-specificity[5]. Combined with fruitful fluoresce proteins[6, 7] and indicators[8], lustrous and remarkable advances in enriched fluorescence microscope[1, 9–12] have brought flourishing discoveries across many disciplines including cell biology[13], immunology[14], and neuroscience[15], among others. However, the limited photon budget with insufficient signal-to-noise ratio (SNR) becomes a fundamental lingering challenge to be solved for fluorescent microscopes that prevents more discoveries to be achieved[16]. The low quantum yield of fluorescent indicators and the stochastic nature of noise make the contamination inevitable[6], aggravating the measurement uncertainty and impairing downstream quantitative analysis, including cell segmentation[17], cell tracking[18], and signal extraction[19]. Overcoming this limitation physically requires enlarging excitation dosage[20] or enriching the expression of indicators[21], but either damaging the fragile living systems or poisoning the cellular health and both altering morphological and functional interpretations that follow. Such a condition is even worse in long-term imaging that necessitates repeated dosage over the same sample hundreds and thousands of times to observe pivotal processes like cell proliferation[22], migration[13, 23], organelle interactions[24, 25], and neuronal firing[26]. To mitigate noise contaminations without excessive light exposure-induced photobleaching and phototoxicity that perturbs the sample in its native state, people have to sacrifice imaging speed, resolution, or dimensions[27].

Despite limited advances achieved across physical approaches, numerous algorithmic approaches have been proposed to break the shot noise limit by utilizing statistics of the noise[28]. Traditional denoising methods that exploit canonical properties of the noise (such as Gaussianity[29] and structures in the signal[30]) achieve great success in photographic denoising [30] but have limited performances in complex, turbulent, and dynamic living systems and with remarkable time consuming and computing complexity. In contrast, supervised learning methods utilizing a data-driven prior learned from paired noisy and clean measurements are proven to be valid as long as samples are drawn from the same distribution[31]. To extend the generalization ability, the requirement of clean data can be further replaced by additional independent noisy measurements[32], fertilizing breakthroughs in interpolating noise-contaminated functional data [33, 34]. However, neither of these supervised methods circumvents the denoising of videographic high-resolution recording with both intensity fluctuations and deformations of living organisms or organelles. The causes of the shortage are manifolds. Firstly, since the same physiological phenomenon would not repeat twice for each cell or organism, the requirement of clean data by methods can only be satisfied through simulations which remain remarkable gaps between training and inferring domains[35]. Secondly, even only the paired noisy data is required in interpolation-based methods like DeepInterpolation [34] and DeepCAD [33], the precondition of interframe continuity likely defiles visualizing rapid transformations of living organisms or organelles. Thirdly, data-hungry nature ensued from the insufficient exploitation of noise statistics forces these methods to compromise, either hampering the genuine visualization to keep the organism safe, or sacrificing the sample health to acquire excessive captures for ensuring visualization quality.

Here, we overcome the aforementioned limitations and propose deep self-supervised learning enhanced microscope (DeepSeMi), a brand-new tool that readily and veritably increases the SNR over 12 dB across various conditions and systems, and catalyzes noise-free videography of diverse structures and functional signals with minimized photodamage in a long term. DeepSeMi explores noise priors that root in data itself through concatenating newly designed eccentric convolution filters and eccentric blind convolution filters with intentionally limited receptive fields across both spatial and temporal dimensions (Supplementary Fig. 1, Methods), fundamentally surrogating the data-hungry shortage of supervised methods and genuinely accomplishing SNR reinforcing even over fast transformed samples. Compared to recently developed interpolation-based denoising methods, DeepSeMi is capable of observing organelles of sophisticated movements and transformations without motion artifacts. Thereby, DeepSeMi standing by the means of self-supervision outperforms other methods in both performance and generalization abilities, and computationally amplifies the photon budget of multiple instruments in long-term tracking of organelles and organisms’ activities without the burden of exacerbating sample health in traditional approaches. Through DeepSeMi, organelle interactions in their native states inside 4-color-labelled-L929 cells were recorded over 30 minutes and 14, 000 time points in high SNR by a confocal microscope, a widely used instrument adored by the high resolution and hated by the photodamage. Aided by DeepSeMi, brittle structures like migrasomes and retractosomes were densely tracked in a half-day-long session uninterrupted without trackable photobleaching, and multiple organelles can be segmented accurately free of false positives by noise contamination. Even fragile and phototoxicity-susceptible samples like *Dictyostelium* cells were also clearly recorded over 36,000 shots in multicolor, attributed to DeepSeMi enhancement.

Not limited to cultured cells and organisms, the capability and generality of DeepSeMi are also demonstrated in a series of photon-limited imaging experiments over various species, including nematodes, zebrafish, and mice, all intravitally. We open-source DeepSeMi to the whole community and hope it can spur new discoveries that were previously unseen by the walls of noise limitation.

## Results

### DeepSeMi roots in noise statistics and accomplishes single-flow high-fidelity denoising

Given the complexity of noisy conditions and sample topologies, limited research has been conveyed to solve the noise contamination of cellular videography. To our knowledge, no data-driven methods capable of long-duration imaging in the intercellular environment at high spatiotemporal resolution have been demonstrated with robust denoising capability in practice. Recent advances in computer vision provide clues to mitigate the problem, where the mapping between different captures of the same scene can form a deep neural network that effectively removes noises in fresh capture[32]. However, such exploitation of noise statistics only stays at the frame level and loses motion information of non-stop contents, limiting applications on spatial-invariant functional imaging or sluggish cell migration in low resolution [33, 34].

The innovation of DeepSeMi roots in a full exploitation of noise statistics. Studies show that mutual mappings from neighbors to a centered pixel can be well established even excluding the pixel itself due to local structure continuity [36]. Under noisy conditions, although those mappings are significantly defiled, the zero means and independence of noise make the average of the defiled mappings relocate the clear pixel information, facilitating estimation of each clear pixel from the surrounding noisy spatiotemporal neighborhood [37] (Fig. 1a). Based on that observation, DeepSeMi thereby established mappings between per pixel of the noisy videography and its surrounding pixels to effectively denoise videography. The utility of pixel-level noise statistics makes DeepSeMi robust even over a single noisy shot, and consequently eliminates the annoying need for excessive captures to ensure the performance compared with previous techniques [33, 34] (Fig. 1e).

**Fig. 1.**
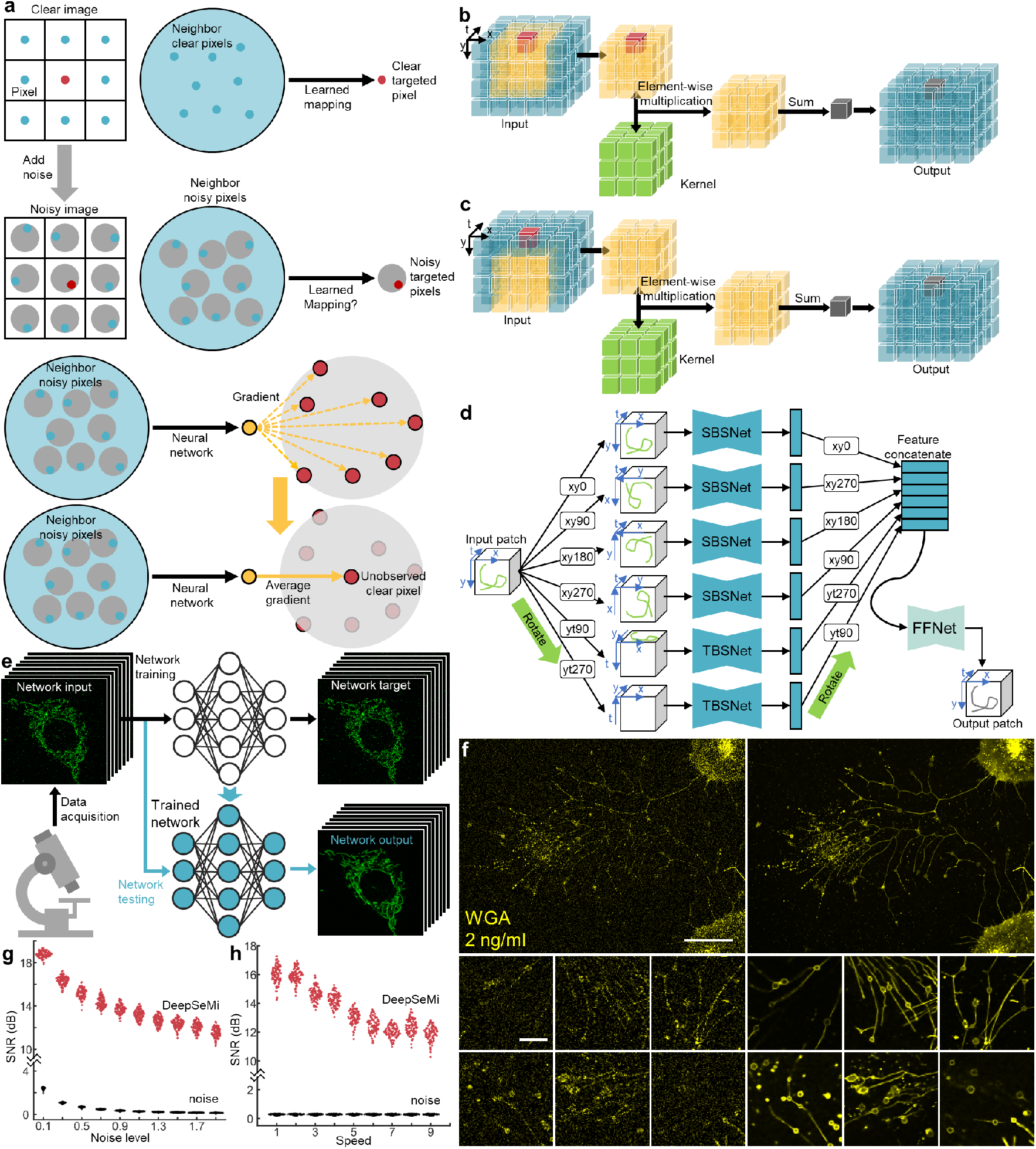
DeepSeMi accomplishes self-supervised video denoising based on the statistical characteristics of noise. **a**, Statistical principle of DeepSeMi. In clean conditions, a learned mapping from neighbors to a centered pixel can be well established even excluding the pixel itself since local structure continuity (the first row). It is worth exploring whether we can establish a learned mapping from neighbor noisy pixels to the targeted noisy pixel.(the second row). A neural network aimed at establishing the learned mapping thereby yields averaged fluctuant gradients on the target pixel (the third row). Fortunately, the zero-mean assumption of noise contaminations ensures that averaged gradients relocate the clean information of the target pixel that is unobserved (the fourth row), which provides the rationale of DeepSeMi. **b**, The schematics of the 3D eccentric convolution. In a 3D (x, y, t) patch (blue), an eccentric neighborhood (yellow) surrounding the target pixel (red) is multiplied with a learnable kernel (green), and the dimension-reduced summation forms an output pixel (grey) in the output patch. Note in eccentric convolution, the eccentric neighborhood still contains the target pixel. **c**, The schematics of the 3D eccentric blind-spot convolution. All symbols are the same as **b**. Note in eccentric blind-spot convolution, the eccentric neighborhood doesn’t contain the target pixel, and thereby the output pixel (gray) excludes the information of the target pixel (red). **d**, Structure of the proposed spatiotemporal hybrid 3D blind-spot convolutional neural network. The neural network consists of six sub-networks with the same structure and a final feature fusion network (FFnet). Among six sub-networks, four spatial 3D blind-spot convolutional neural networks (SBSnet, top four) and two temporal 3D blind-spot convolutional neural networks (TBSnet, bottom two) share the same parameters, respectively. The input patch is rotated and fed into each sub-network, and the output features accordingly are rotated in order to match each other’s size before feature fusion (Methods). **e**, DeepSeMi enables signal-to-noise-ratio (SNR) enhancement with only the experimental data through a single shot. The low-SNR recordings can be used to train the proposed self-supervised neural network *in situ*, which enable the trained network to enhance the low SNR recordings itself. **f**, The raw captures of cells labeled by WGA with a diluted concentration (2 ng/ml, 500 times diluted, left) and corresponding DeepSeMi enhancement (right). Magnified views about migrasomes in 500 times diluted dye concentration are presented at the bottom (left for raw and right for DeepSeMi enhancement). Scale bar in the first row is 30 μm, and in the second rows is 10 μm. **g-h**, DeepSeMi denoising performance indicated by the SNR over different noise levels (Supplementary Fig. 3) and content speeds (Supplementary Fig. 4), respectively.

To establish these special mappings, two brand new convolution kernels were developed for conveying the aforementioned thought with optimized efficiency in DeepSeMi. The first convolutional kernels receive both the inferred pixel and its eccentrically surrounded neighbors to keep the DeepSeMi efficient in both restoring structures and eliminating noise (Fig. 1b, Supplementary Fig. 1b), and are accordingly named as eccentric convolution. The second convolution kernels resemble the blind-spot property by only receiving the eccentrically surrounded neighbors of the inferred pixel to achieve an even stronger noise cleanse ability (Fig. 1c, Supplementary Fig. 1c), and are accordingly named as eccentric blind-spot convolution. A single flow across the blind-spot convolution thereby consists of millions of sub-approaches where each input noisy pixel is synthesized only by the neighbors without itself, accomplishing denoising in a self-supervised learning manner exceedingly efficiently. The rationale for combining both filters in the DeepSeMi is to achieve an appropriate balance between the preservation of details and noise robustness with the assistance of the pixel-level blind-spot technique (Methods). Six branches composed of these two convolutional filters deliver permutational receptive-limited fields of both spatial and temporal dimensions, and are further merged by a feature fusion network to form preferential representations of the output video block (Fig. 1d). Computation losses are differentiated ergo between the input and output to guide the updates of the network parameters through backpropagation (Supplementary Fig. 2). The comprehensively optimized DeepSeMi also leverages a time-to-feature folding operation which feeds more temporal information without increasing additional computational cost to increase performance (Methods).

We benchmarked the denoising capability of DeepSeMi through extensive simulations compared with various mainstream methods. To fully emulate real experiments within complex situations, we evaluated those methods in Moving MNIST datasets where both the noise level and the movement speed of the contents are varied in a large range. Among all methods, DeepSeMi achieved the best denoising results across all noise levels, even achieving 15 dB higher SNR compared to raw capture at extremely noisy conditions where samples were submersed in noise (Fig. 1g, Supplementary Fig. 3). While most of the literature merely comparing SNR in static scenes, we further evaluated the denoising ability of those methods encountering swift contents across various speeds. As the increase of the content moving speed, DeepSeMi kept being the top tier in terms of restoration quality over other methods with at least 12 dB SNR improvement (Fig. 1h, Supplementary Fig. 4), where techniques using frame-level noise statistics (DeepCAD [33] and DeepInterpolation [34]) lowered their performance quickly due to the frame interpolation nature (Supplementary Fig. 5, Supplementary Fig. 6). In more complicated Poisson noise contaminations where the noise scale correlates with the image intensity (Supplementary Fig. 7), DeepSeMi still outperformed all other methods by over 4 dB ahead. Across all tests, the UDVD15 technique [38] utilized the similar blind-spot technique immediately following up DeepSeMi. However, our proposed DeepSeMi achieved superior performance thanks to the improvement of spatiotemporal convolutions, additional eccentric blind-spot convolutions, and additional receptive field limited branches in temporal domain (Methods).

Besides, DeepSeMi was also proved to have generalization ability across different noise scales and content speed. Given the DeepSeMi that trained at a moderate speed (Supplementary Fig. 8a-c), the performances are nearly consistent when the content speed varies across 20% to 180% range. We further tested the generalization ability of DeepSeMi in experiments, where DeepSeMi was trained for the modality of mitochondrial membrane but tested in the co-labeled cell membrane and mitochondrial matrix data (Supplementary Fig. 9a). We found the noise-contaminated mitochondrial matrices were cleaned by DeepSeMi in both clustered forms close to the cell center and scattered forms in the cell edge (Supplementary Fig. 9b-e). Composited interactions of both membranes and mitochondrial matrix were clearly displayed after DeepSeMi enhancement which was only trained in a third and unimodal data (Supplementary Fig. 9f-h). By denoised dual-color co-labeled mitochondria data (Supplementary Fig. 10a), self-consistency of DeepSeMi was validated since the denoised results were highly consistent between dual channels despite the noise distributions being largely different between them (Supplementary Fig. 10c). The great generalization ability and self-consistency of DeepSeMi ensure the fidelity of observation across complicated micro-environment during long-term cellular imaging, accomplishing apparent enhancements in recovering both structural and functional diversities (Supplementary Fig. 11, Supplementary Video 1).

### DeepSeMi unlocks high-speed long-term imaging with minimized photobleaching

High-temporal resolution imaging is ideal for observing swift intracellular organelle interactions, cell migration, and multicellular interactions, yet regretfully limited in a short term due to the compromise of photobleaching and phototoxicity. With extensive evaluations, we found that healthy mitochondria can only stand for 45.3 μW laser power (2%, 488 nm) (Supplementary Fig. 12) for a 3-minute-long session at 30 frames per second (fps) in a commercial confocal microscope without apparent photobleaching (Supplementary Fig. 13, Methods). Higher scanning laser dosage quickly quenched the fluorescence, failing the imaging process due to missing mitochondrial structural information. However, such a low power dosage exacerbated the noise contaminations to the observations and yielded barely characterized structures (Supplementary Fig. 13d), and the situation was even worse when the mitochondria were densely clustered due to lack of sparsity. On the other hand, with the proposed DeepSeMi, mitochondria under even 14.6 μW (0.5%, 488 nm) power dosage can be faithfully denoised with intact and natural form restored (Supplementary Fig. 14, 15). Under that mild excitation, the fluorescent intensity drop was unrecognizable, suggesting DeepSeMi enhancement not only accomplished high-temporal resolution recording but even reduced the photobleaching further (Supplementary Video. 2). From other perspectives, the computational enhancement of DeepSeMi brings a surge of available photon budget of optical instruments. Considering DeepSeMi achieves even higher visualization quality of mitochondrial structures in 23.1 μW (1%, 488 nm) (Supplementary Fig. 13b) than raw captures in 537 μW (32%, 488 nm) (Supplementary Fig. 13g), the available photon budget was enlarged at least ten folds.

We quantitatively verify the photon budget enlargement of DeepSeMi across two dimensions. In the first dimension, we approximated the photon budget enlargement as the multiplication of excitation power in raw captures through which the same SNR of DeepSeMi enhancement can be achieved (Supplementary Fig. 16). We found at least 15-times more power dosage in raw frames was required to produce the same level of imaging quality as DeepSeMi enhancement across various noisy conditions, verifying DeepSeMi enlarges the photon budget by 15 folds leastways. In the second dimension, we investigate the photon budget enlargement as the excessive concentration of dyes in raw captures to approach the DeepSeMi-enhanced SNR. We proved DeepSeMi achieved no-compromise results in over 50 times diluted dye concentrations across migrasomes, lysosomes, and mitochondria, and the resulting captures were comparable with the non-diluted ones (Fig. 1f, Supplementary Fig. 17). Although both the higher power dosage and dye concentration facilitate better visual inspections with fewer noise contaminations, on the other hand, both of them cause significant cytotoxicity and perturbation over the native regulation of the organelles and organisms. Instead, DeepSeMi enables tens of times photon budget increments computationally, permitting high-fidelity functional and structural interrogation which is previously unmet. Towards directions of broader applications, the multiplied photon budget by DeepSeMi strongly extends the capacity of the commercial confocal microscope in pursuing higher spectral complexity, higher frame rate, and longer recording sessions.

With the encouragement of apparent SNR enhancement of DeepSeMi under sample-friendly power dosage across thousands of captures, we performed imaging at 7.5 fps on L929 cells with four structures labeled by four colors (tagBFP-SKL, TOM20-GFP, SiT-mApple, and WGA647 for peroxisomes, mitochondria, Golgi, and migrasomes, respectively) on a commercial confocal microscope (Fig. 2a, Methods), for 30 minutes and over 13,500 time points. Excitation power was set at 2% to obviate photobleaching and keep live cells healthy (Fig. 2b), at the expense of plenty of noise and ruptured structures that defiled the raw captures. Contrastingly, the enhancement of DeepSeMi clearly revealed delicate structures of punctate peroxisomes, threadlike mitochondria, and fluctuated membranes (Supplementary Video. 3). The brittle mitochondrial fission and fusion were obviously distinguished (Fig. 2c-d), highlighting the importance of combining minimization of illumination photon dose with SNR enhancement of DeepSeMi.

**Fig. 2.**
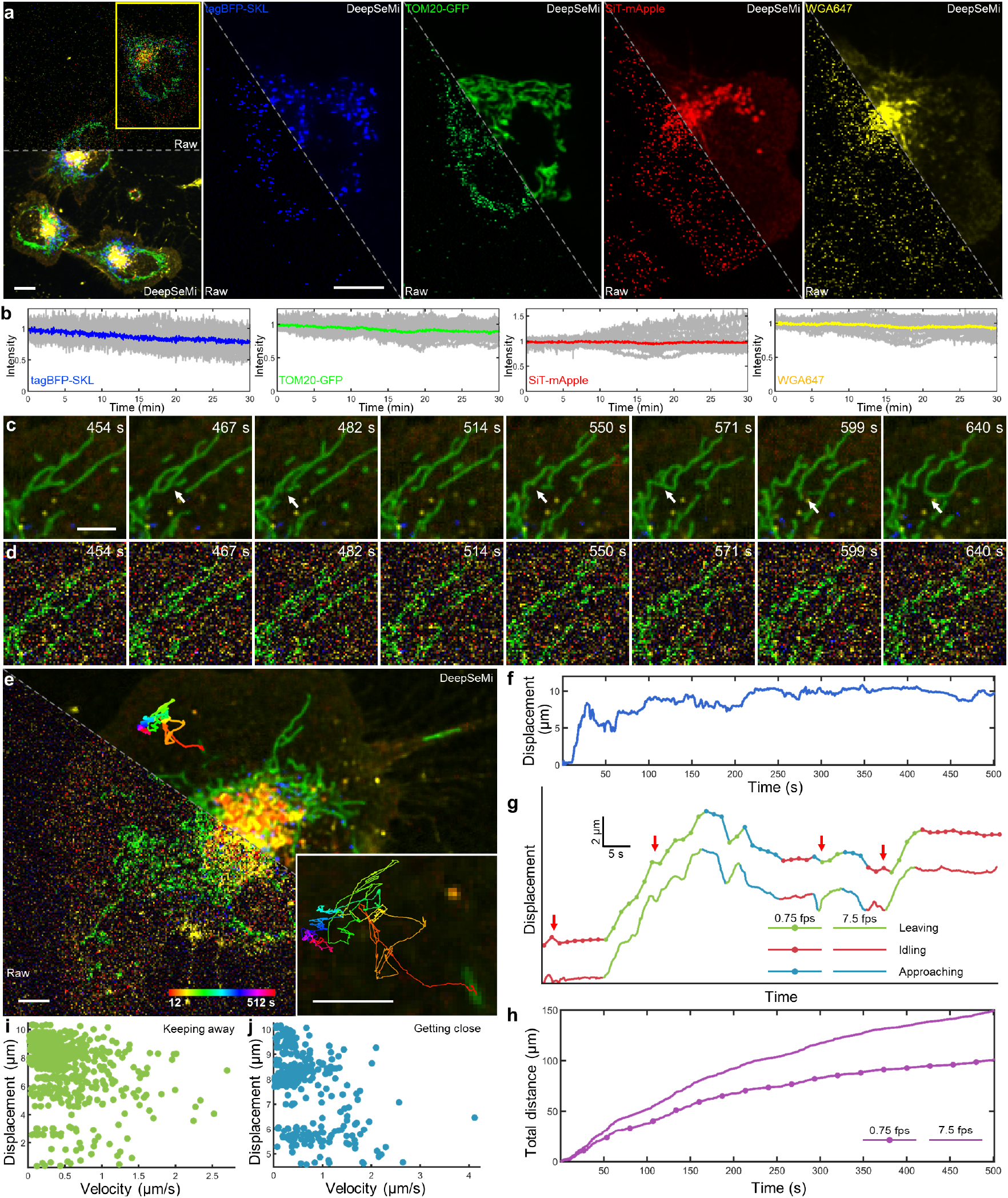
Long-term, high-temporal resolution, and low-phototoxicity imaging of organelle interactions by DeepSeMi. **a**, Left, raw (top) and DeepSeMi-enhanced (bottom) micrographs of an L929 cell expressing fluorescent proteins (TOM20–GFP, TagBFP–SKL, and SiT-mApple) and labeled by WGA647. Right, individual channels of the yellow box marked in the left panel are displayed separately. Scale bar, 10 μm. **b**, Fluorescence intensity fluctuations *(n* =10) of 4 channels during a 30-minute long imaging session (13,500 frames) at 2% light intensity. Fluorescence intensity curves were normalized to initial values. **c-d**, Raw and DeepSeMi-enhanced time-lapse images that reflect mitochondrial morphological changes during low-light recording. White arrows mark the process of mitochondrial fission and fusion. Scale bar, 5 μm. **e**, Raw (left) and DeepSeMi-enhanced (right) four-color cellular imaging in low-light conditions, with trajectories of a rod-shaped mitochondria tracked and zoomed in the bottom-right corner. The color-coded time stamps are labeled at the bottom. Scale bar, 5 μm. **f**, Displacement of the rod-shaped mitochondria plotted as a function of time. **g**, Inferred mitochondria displacements versus time under different imaging frame rates. Different colors represent different relative states of rod-shaped mitochondria to the cell body, where green for leaving, blue for approaching, and red for idling. Red arrows mark differences between displacement inferences of full sampling rate (7.5 Hz) and 10-fold sub-sampling rate (0.75 Hz). **h**, Tracked drifting distances of mitochondria during 500 seconds with full sampling rate (7.5 Hz) and 10-fold sub-sampling rate (0.75 Hz). **i-j**, Distributions of the moving rates and displacements of tracked rod-shaped mitochondria during leaving and approaching states, respectively.

Together with the high temporal resolution and long-term capability, DeepSeMi catalyzes new abilities of tracking subtle movements of mitochondria, an important component of mitochondria regulation in many aspects of cell biology. An individual rod-shaped mitochondrion was tracked based on DeepSeMi-enhanced recordings over 500 seconds, unveiling complicated trajectories and nonlinear movements (Fig. 2e-f). Sampling the data at full temporal resolution presented brief transitions between mitochondria leaving and approaching, and quick motions happened when the leaving or approaching of mitochondria paused temporally [39] (Fig. 2g). These transient processes cannot be captured if the sampling frequency dropped by 10-fold to 0.75 Hz, which was the compromised framerate for the standard confocal microscope without DeepSeMi enhancement in catching the similar photon budget. We thereby demonstrated that the high temporal resolution enabled by DeepSeMi is indispensable to characterizing the veritable trajectories as complex movements between frames were likely to be missed when temporal resolution dropped down (Fig. 2h). We measured mitochondria leaving and approaching rates of 0.53 μm/s and 0.46 μm/s, respectively. Furthermore, when analyzing these rates as a function of the displacement of each leaving or approaching event (Fig. 2i-j), we found that long displacing events correlated with slow rates of leaving or approaching. There was a broader range of leaving rates compared to approaching rates during short displacing events, leading to diverse fluctuations in mitochondria displacement. Overall, the SNR enhancement of DeepSeMi vehemently enlarged the available photon budget of an optical instrument without compromising visual quality for down-stream analysis. DeepSeMi allowed us to quantify not only mitochondria dynamic displacements but also alterations of other organelles on a much finer temporal scale than what was achieved in previous methods.

### DeepSeMi enables monitoring migrasomes and retractosomes over a half day in their native states

Migrasome is recently recognized as an extracellular organelle that plays a significant role in various physiological processes, including mitochondrial quality control, organ morphogenesis, and cell interaction [40, 41]. Despite fruitful results that have been discovered related to migrasome regulations by light microscope, uninterruptedly observing migrasomes during cell migrations in a half-day-long term remains challenging limited by continuously imaging-induced photobleaching and phototoxicity (Supplementary Fig. 18).

Here, through DeepSeMi enhancement, we accomplished high-resolution 2 fps imaging of the generation, growth, and rupture of migrasomes in a half-day-long term with over 86,000 time points with only 2% power shots (45.3 μW of 488 nm, 49.8 μW of 561 nm). L929 cells expressed TOM20-GFP and TSPAN4-mCherry to tag the mitochondria and migrasomes, respectively. A representative two-color image frame from a movie of the mitochondria and migrasomes clearly showed the enormous SNR enhancement by DeepSeMi compared to the raw capture (Fig. 3a, Supplementary Video 4). Near the cell body, DeepSeMi enabled us to find migrasomes that presented the entire generation and growth procedure across ~ 300 minutes of imaging windows, which was 41% of the whole imaging session (Fig. 3b). The DeepSeMi enhanced results clearly show that some mitochondria were expelled by the cell and kept inside a migrasomes (Fig. 3d-e), known as the mitocytosis [41]. Compared to barely recognized migrasomes in the raw images (Fig. 3c), 51 migrasomes were segmented from the whole DeepSeMi-enhanced capture (Methods), with color-coded area and longevity statistics summarized in Fig. 3f. We measured an averaged maximum area of 5.81 μm^2^ (Fig. 3g) during an averaged 141-minute lifespan of migrasomes (Fig. 3h), which were weakly correlated with each other (Fig. 3i). We noticed a general pattern of the maximum area across those migrasomes consisting of a quick rising representing the growth, a slightly declined plateau, and a sharp drop representing the rupture (Fig. 3j). The dynamics of rupture was much faster than the other two procedures (Fig. 3k), which necessitated DeepSeMi enabled high temporal-resolution and uninterrupted captures across a long term to catch these features.

**Fig. 3.**
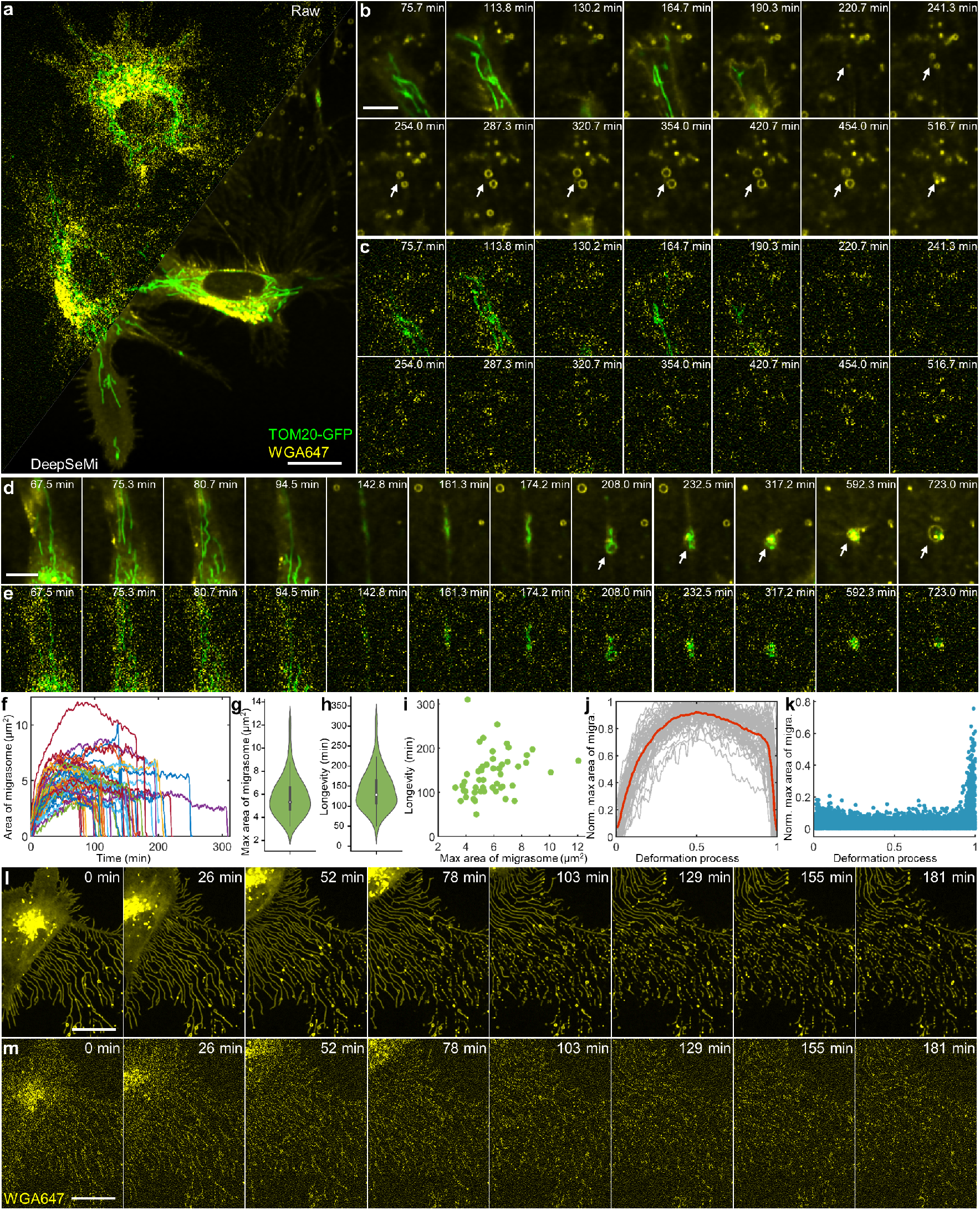
DeepSeMi enables half-day-long observations of migrasomes and retractosomes with low-phototoxicity. **a**, Raw (top left) and DeepSeMi-enhanced (bottom right) micrograph of L929 cells expressing both TOM20-GFP and TSPAN4-mCherry. Scale bar, 20 μm. **b-c**, Zoom-in panels that visualize extracellular migrasomes generation and displacement by raw and DeepSeMi-enhanced recordings, respectively. The migrasomes marked by white arrows burst at the end of their longevities. Scale bar, 10 μm. **d-e**, Zoom-in panels that visualize mitocytosis and displacements by raw and DeepSeMi enhanced recordings, respectively. Scale bar, 10 μm. **f**, The areas of extracellular migrasomes changing along time in DeepSeMi-enhanced videos. Different colors represent different migrasomes (*n* = 51). **g**, Violin plot of the maximum area of extracellular migrasomes in DeepSeMi-enhanced videos. White circle: median. Thin vertical lines: upper and lower proximal values. Violin-shaped area: kernel density estimates of data distribution. *n* = 51 data points. **h**, Violin plot of the longevity of extracellular migrasomes in DeepSeMi-enhanced videos. Symbols are the same as in **g**. *n* = 51 data points. **i**, Scatter plot of longevity and maximum area of extracellular migrasomes in DeepSeMi-enhanced videos. *n* = 51 data points. **j**, Statistics of the normalized migrasome area changing across the migrasomes life span. Gray curves represent the trend of each migrasome (*n* = 51), and the red curve represents the average. **k**, Histogram of the area changing rate of migrasomes across *n* = 51 migrasomes. **l-m**, Generation of retractosomes in regions through which cells have migrated over. A global view where the first row represents images enhanced by DeepSeMi and the second row represents the raw images. Scale bar, 20 μm.

Retractosome is recently reported as a newly discovered extracellular organelle that is closely related to cell migrations [42]. Since uninterrupted cell migrations can be continuously imaged benefiting from DeepSeMi-enabled low-phototoxicity, high-SNR, and long-term recording ability, pronounced retractosomes were recognized which were transformed from broken-off retraction fibers (Fig. 3l-m). Albeit the indistinguishable beads-on-a-string features in the raw captured video, retractosomes were clearly recognized when they moved along with the wobbled retraction fibers (Supplementary Video 5). After the cell migrated away, plenty of retraction fibers and retractosomes were left behind and forming a complicated network structure that was fractured by the noise. In opposition, DeepSeMi reunited the network by wiping out noise contaminations and thus delivering the potential to study the physiological functions of retractosomes in the future.

### DeepSeMi facilitates automated analysis of cellular structures from massive data

Uncovering the peculiarities of important life-preserving and disease-driving organelles requires robust and unbiased segmentation and tracking tools. Compared to biased and time-consuming manual analysis, recent advances in deep-learning-based processing techniques utilize statistical and graphical knowledge to accomplish fast, unbiased, and automated organelle analysis and are capable of recognizing complicated dynamics like fission and fusion of mitochondria [17]. Confronting the growing requirement of long-term recordings and attendant considerable amounts of cellular imaging data in hundreds of gigabytes [43], automated cellular analysis gradually becomes indispensable for new physiological discoveries.

Inspired by those advancements, we utilized the state-of-the-art organelle segmentation method [44] and accordingly trained 3 segmentation networks for mitochondria, migrasomes, and retraction fiber, respectively (Fig. 4a, Methods). We found raw captures of mitochondria under 14.6 μW (0.5% of 488 nm), a bio-friendly power dosage, suffered pronounced segmentation errors due to noise contaminations (Fig. 4b-d, Supplementary Fig. 19). The incorrect segmentation fragments in the background were only eliminated when the power dosage was pushed into 537.6 μW (32% of 488 nm), at a cost of significant photobleaching (Fig. 4b-d, Supplementary Fig. 13h). By contrast, DeepSeMi enhancement enabled the segmentation model to output reasonable and gap-less results even at 14.6 μW (0.5% of 488 nm) (Fig. 4b, Supplementary Fig. 19), permitting reliable segmentation during long-term imaging thanks to heavily reduced photobleaching. Through additionally performing mitochondrial skeletonization and keypoint detection based on instance segmentation[17] (Supplementary Fig. 20), we found remarkable noisy stains in raw captures were recognized as endpoints and junctions of broken skeletons (third row of Fig. 4b, Supplementary Fig. 19). These false positives were well avoided in DeepSeMi enhanced results, and the skeletonization result by DeepSeMi at 14.6 μW (0.5% of 488 nm) are comparable of that in the raw image at 537.6 μW (32% of 488 nm). Quantitively, DeepSeMi enhanced videography achieved significantly larger mitochondria area (Fig. 4e, ***p<0.0001, two-sided Wilcoxon rank sum test; Supplementary Fig. 19, Methods) and longer branch length (Fig. 4f, ***p<0.0001, two-sided Wilcoxon rank sum test; Supplementary Fig. 19, Methods) compared to the raw ones at sample-friendly power dosage (14.6 μW (0.5% of 488 nm) and 23.1 μW (1% of 488 nm)). These statistics were only comparable when the power comes to harmful 537.6 μW (32% of 488 nm, p>0.1, two-sided Wilcoxon rank sum test). The over 15 times power reduction of DeepSeMi in achieving high-quality subcellular segmentation validated with over 15 times enlarged photon budget in photobleaching study previously (Supplementary Fig. 13), together indicate the strong promotion of DeepSeMi over an optical instrument in terms of bio-friendly, resolving ability, and data fidelity.

**Fig. 4.**
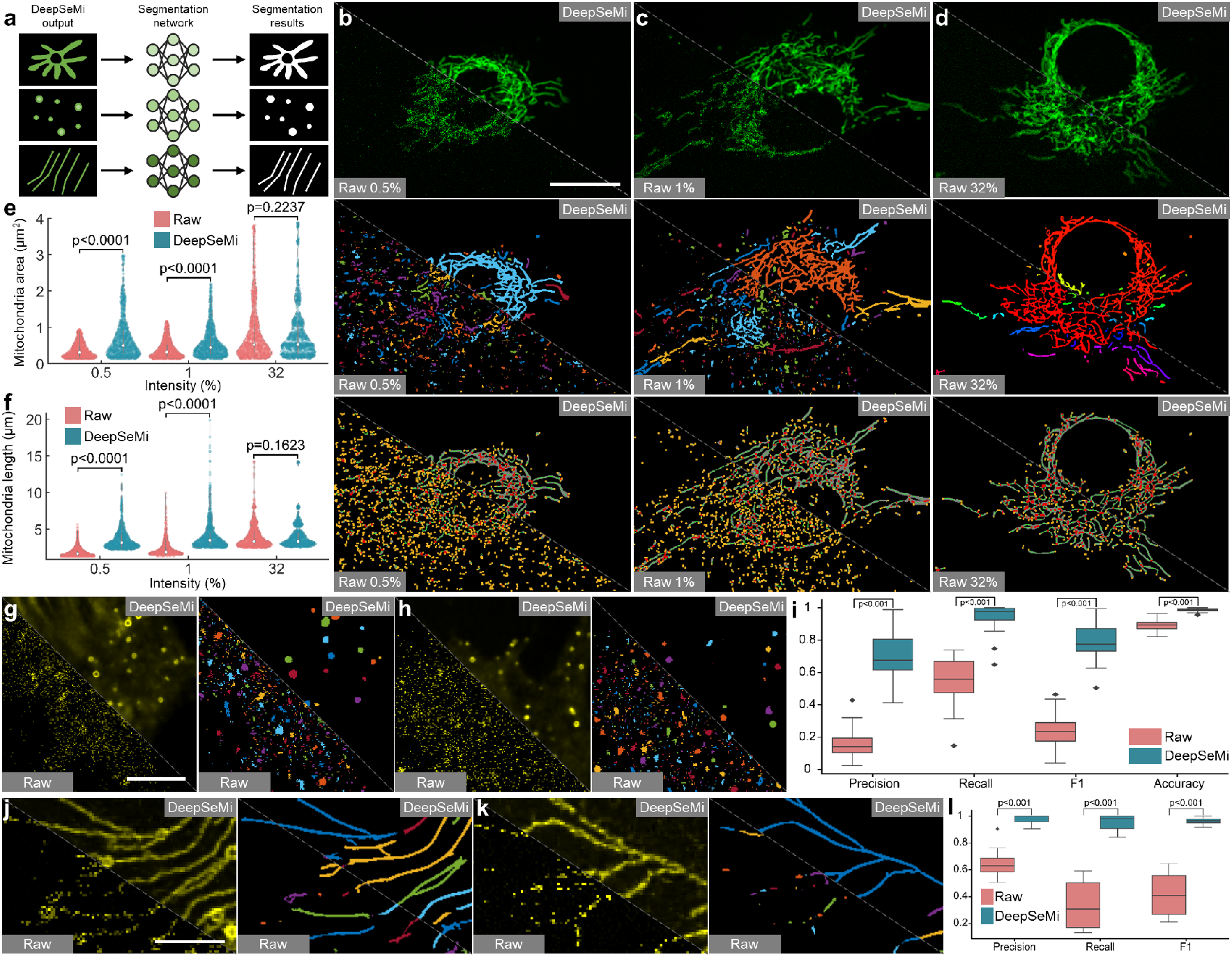
DeepSeMi facilitates accurate automated analysis of cellular structures with low light dosage. **a,** Schematic diagram illustrating the segmentation of mitochondria, migrasomes, and retraction fibers through three neural networks (Methods). **b-d**, Differences of mitochondrial analysis based on raw images (bottom left) and DeepSeMi-enhanced (top right) images decreased as power dosage increased. The first row represents the raw captures (bottom left) and the DeepSeMi-enhanced fluorescence images (top right). The second row represents the instance segmentation of the raw captures (bottom left) and the enhanced images (top right). The third row represents the skeletonization of the raw captured mitochondria (bottom left) and the enhanced mitochondria (top right). Scale bar, 20 μm. **e**, Statistics of mitochondria area based on the instance segmentation before (red) and after DeepSeMi enhancement (blue). White circle: median. Thin vertical lines: upper and lower proximal values. Violin-shaped area: kernel density estimates of data distribution. **f**, Statistics of branch length of mitochondria based on the skeletonization before (red) and after DeepSeMi enhancement (blue). Symbols as in **e**. **g-h**, Instance segmentation of migrasomes before (bottom left) and after DeepSeMi enhancement (top right). Scale bar, 20 μm. **i**, Segmentation precision, recall, F1, accuracy scores of the migrasomes before (red) and after DeepSeMi enhancement (blue). Ground truth data is manually annotated (Methods). *n* = 32 images. **j-k**, Instance segmentation of retraction fibers before (bottom left) and after DeepSeMi enhancement (top right). Scale bar, 10 μm. **l**, Segmentation precision, recall, F1 scores of the retraction fibers before (red) and after DeepSeMi enhancement (blue). Ground truth data is manually annotated (Methods). *n* = 12 images.

To further evaluate the improvement of segmentation accuracy brought by the DeepSeMi enhancement, we manually segmented migrasomes and retraction fibers as the ground truth and then compared the results with automated segmentations on DeepSeMi enhanced videography (Methods). DeepSeMi apparently achieved much clearer micrographs and hence cleaner segmentations (Fig. 4g-h). Statistically, DeepSeMi enhancement achieved 0.9449 ± 0.0782 recalls (*n* = 32 images) in migrasome segmentations, holding a safe head compared to raw-video-based segmentation (0.5522 ± 0.1359 recalls, *n* = 32 images). The same advantages were held in segmenting string-like retraction fibers (Fig. 4j-k), where DeepSeMi enhancement achieved 0.9493 ± 0.0618 recalls (*n* = 12 images) compared to 0.3391 ± 0.1848 recalls by raw video (*n* = 12 images, Fig. 4l). The high segmentation accuracy enabled by DeepSeMi under sample-friendly power dosage would be the key for massive data analysis through automated algorithms after long-term recordings.

### DeepSeMi accomplishes SNR enhancement across various samples

Lastly, we demonstrated that DeepSeMi effectively increases SNRs across various samples, including cultured cells, unicellular organisms, nematodes, non-mammalian vertebrates, and mammals. We have demonstrated DeepSeMi enabled high-temporal-resolution imaging of mitochondria, low-phototoxicity half-day-long imaging of migrasomes and retractosomes, and facilitated automated analysis in massive data under bio-friendly illumination dosage, but the power of DeepSeMi could be extended further. Here we delineated DeepSeMi helped study of rearrangement of organelles after decomposing cytoskeleton and other organelle-related studies. By dosing an appropriate concentration of latrunculin-A (lat-A) to induce the depolymerization of the intracellular cytoskeleton, a new spatial distribution of intracellular organelles was formed (Supplementary Fig. 21). We found the migrasomes were generated following the rapid contraction of the cell membrane after depolymerization of the cytoskeleton (Fig. 5a). All those observations relied on the enhancement of DeepSeMi, which restored mitochondria and other organelles of diverse morphologies from noise. Similar improvements happened in the study of vesicle fission (Supplementary Fig. 11h, Supplementary Video 1), where kymographs (x-t projections) obviously presented the enhancements of DeepSeMi (Supplementary Fig. 11i), and also in the study of migrating cell interacting with a migrasome (Supplementary Fig. 22b), producing migrasomes (Supplementary Fig. 22c), and expelling mitochondria in low light dosage (Supplementary Fig. 22d, Supplementary Video 6).

**Fig. 5.**
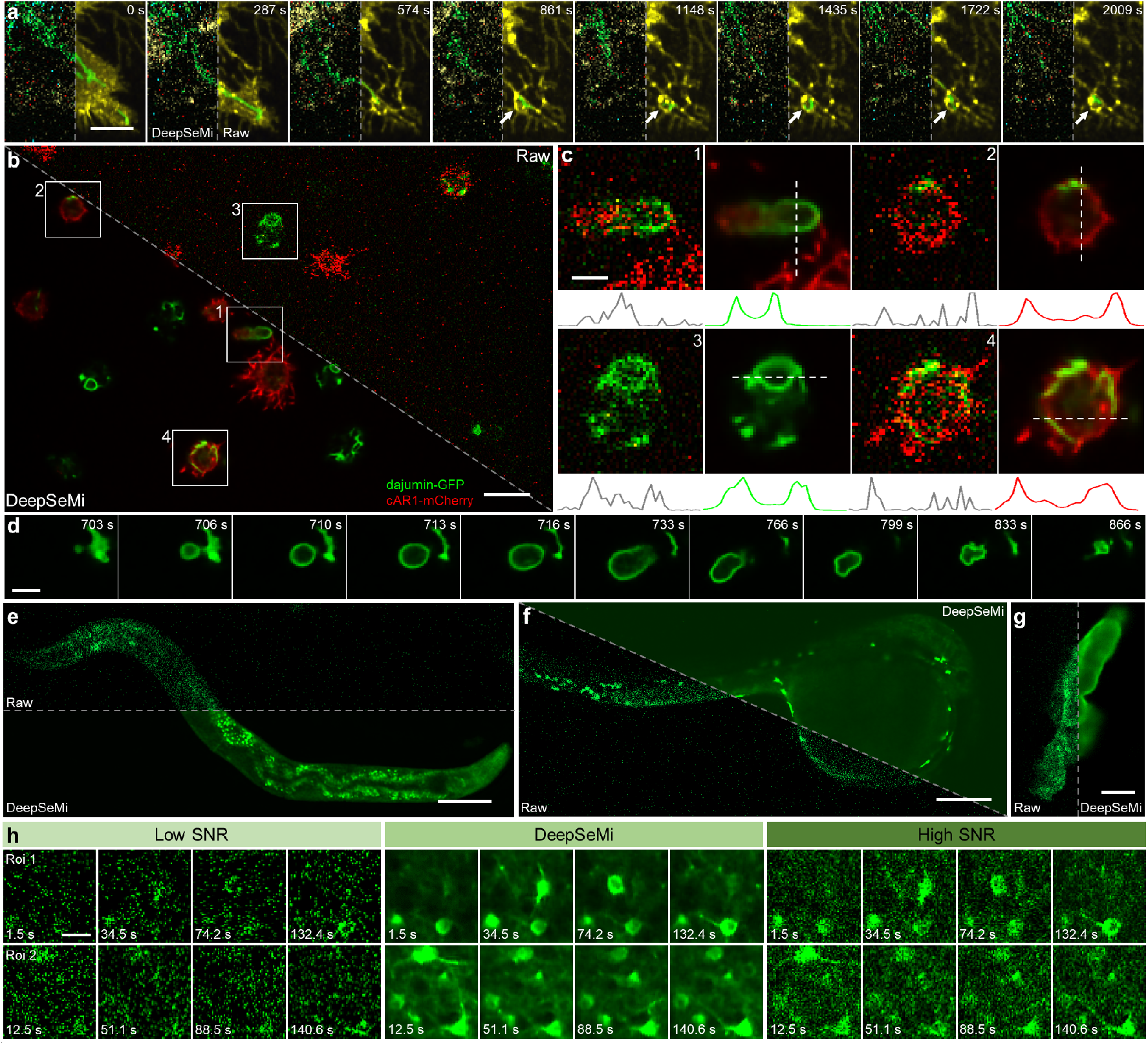
DeepSeMi seamlessly improves SNRs over various species. **a**, Generation of a migrasome from the L929 cell with four organelles labeled colorfully (TOM20–GFP, WGA647, TagBFP–SKL, and SiT-mApple, Supplementary Fig. 20) after treatment with Latrunculin-A (lat-A) (0.5 μg/ml. Methods). For each panel, the right part represents DeepSeMi-enhanced results and the left panel represents the raw image. Scale bar, 10 μm. **b**, Raw (top right) and DeepSeMi-enhanced (bottom left) long-term high-speed imaging of photo-sensitive *Dictyostelium* cells. Scale bar, 10 μm. **c**, Zoom-in panels of the white boxes marked in **b** represent contractile vacuoles and membranes. Intensity profiles along the white dash lines were plotted at the bottom. Scale bar, 3 μm. **d**, Time-lapse imaging of expansion and contraction of the contractile vacuole enhanced by DeepSeMi. Scale bar, 4 μm. **e**, *In vivo* imaging of *C. elegans* in a millimeter-scale field-of-view by raw (top) and DeepSeMi-enhanced (bottom) captures, respectively. Scale bar, 100 μm. **f**, *In vivo* imaging of zebrafish larvae in a millimeter-scale field-of-view by raw (top right) and DeepSeMi-enhanced (bottom left) captures, respectively. Scale bar, 200 μm. **g**, Observation of macrophage in zebrafish larvae *in vivo* by raw (left) and DeepSeMi-enhanced (right) images, respectively. Scale bar, 5 μm. **h**, Low-SNR image (left), DeepSeMi-restored image (middle), and high-SNR reference image recorded by 10-fold higher photon flux as references (right). Low-SNR and high-SNR images were recorded through a hybrid multi-SNR two-photon system for validation [33]. 8 time points were displayed for each modality. Scale bar, 20 μm.

DeepSeMi also enabled high-SNR, half-hour-long imaging of *Dictyostelium* cells, an amoeba-like important eukaryote model for studying genetics, cell biology, and biochemistry [45]. Despite the great value of *Dictyostelium* cells in research, it is ultrasensitive to photodamage since 215 μW of laser dosage at 638 nm and 50.6 μW of laser dosage at 561 nm killed 30% of *D. discoideum* after 30-minute imaging, preventing high-SNR half-hour-long imaging in conventional approaches (Supplementary Fig. 23, 24). We applied DeepSeMi to circumvent the problem, which enabled dual-color and high-SNR imaging at the 45.3 μW dosage at 488 nm and the 49.8 μW dosage at 561 nm over 30 minutes without apparent photodamage (Fig. 5b, Supplementary Fig. 23, 25). Contractile vacuoles and membranes of *Dictyostelium* cells were evidently recognized with clear boundaries through DeepSeMi enhancement (Fig. 5c), and uninterrupted videography dedicatedly enabled by DeepSeMi unveiling startling motions of *Dictyostelium* cells such as contracting (Fig. 5d, Supplementary Video 7). The ability of DeepSeMi that strongly improves SNR without increasing power dosage sheds new light on studying photodamage-sensitive but valuable animal models like *Dictyostelium* cells.

*Caenorhabditis elegans (C. elegans)* and zebrafish are used as central model systems across biological disciplines[46, 47]. Rather scattered tissues of *C. elegans* exuberate the noise contaminations even further compared to cultured cells (Fig. 5e, Supplementary Fig. 26a), but DeepSeMi still substantially improved the contrast and sharpness of cells (Supplementary Fig. 26b-f). Although utilizing a higher NA objective suffers even more from scattering, DeepSeMi restored delicate structures with sharp edges and high contrast from noise (Supplementary Fig. 26g-j). On the other hand, the transparency of zebrafish larvae not only helps better observation of structures and functions of cells and organisms *in vivo,* but also eliminates the protective barrier to photodamage during optical observation [48]. Thereby, imaging zebrafish larvae necessitates low illumination power to not alter the sample health state and normal physiological regulation, which inevitably raises challenges from noise contaminations (Fig. 5f, Supplementary Fig. 27a). We proved that enhancement of DeepSeMi broke the dilemma and provided a clear view of macrophage in zebrafish larvae under a mild power dosage (45.3 μW, Fig. 5g, Supplementary Fig. 27b-c), supplying the potential for long-term observation for studying development and function in the highly complex vertebrate model system.

DeepSeMi is also demonstrated to be operative in functional imaging in mice that are widely used in systems and evolutionary neuroscience. We tested the generalization ability of DeepSeMi in nonlinear microscope where neurons were sequentially excited by a focused femtosecond laser *in vivo.* DeepSeMi readily enhanced visualization of morphologies of neuronal structures (Fig. 5h, Supplementary Fig. 28a-c, Supplementary Fig. 29a-i) from barely recognized noisy captures, and also veritably increased temporal contrast of calcium transients (Supplementary Fig. 28d, Supplementary Fig. 29j). The denoised videos by DeepSeMi facilitated 1.5 times more neurons to be found and would impel potential interrogation of neuronal circuits (Supplementary Fig. 28e, Supplementary Fig. 29k). For observing even smaller structures like wobbled neuronal dendrites and axons *in vivo* in the mouse brain, the enhancement of DeepSeMi also has no compromise (Supplementary Fig. 30).

## Discussion

Many species of great scientific value are vulnerable to photodamage, necessitating low-power dosage for sample health yet sacrificing SNR, and the condition deteriorates when high spatiotemporal resolution is required for deciphering composited morphology-related regulations. To address these problems, we present DeepSeMi, a versatile self-supervised paradigm capable of enhancing over 12 dB SNR, improving 15-fold photon budget, and reducing 50 times fluorescent dye concentration across various species and instruments with only noisy images required. DeepSeMi with specially designed receptive field-limited convolutional filters readily accomplishes noise contamination removal without clean data reference or inter-frame interpolations, achieving superior performance over other methods especially in data with complicated transformation. Computationally enhanced photon budget by DeepSeMi fertilized high-frame-rate 4-color organelle recordings across tens of thousands of frames, tracking migrasomes and retractosomes over a half day, and ultra-photodamage-sensitive *D. discoideum* imaging over thousands of frames, all high-fidelity, intravitally, and sample-friendly. Besides, DeepSeMi was proven to help automated analysis of cells and organelles which is a strong aid in processing massive imaging data and is in trend. Performance of DeepSeMi on various species including nematodes, zebrafish, and mice on both widefield and two-photon microscopes was also validated both qualitatively and quantitatively. In conclusion, DeepSeMi offers a combination of high-resolution, high-speed, multi-color imaging and low photobleaching and phototoxicity that makes it well suited for studying intracellular dynamics and beyond.

As a fundamental limitation in fluorescence imaging, stochastic noise determines the upper bound of imaging quality and compromises speed, resolution, and sample health across any instrument. The proposed DeepSeMi can be seamlessly extended to various devices that suffer the noise most, including three-photon microscope with ultra-small absorption cross-section[49], and Raman microscope with critical excitation conditions[50]. In other devices such as widefield and light-field microscope where background contaminates more in scattering tissues than noise, DeepSeMi can collaborate with computational background elimination methods [51] to jointly improve imaging quality with backgrounds rejected and SNR increased.

The rearrangement of computationally multiplied photon budgets by DeepSeMi can be more diverse. We have shown benefits of shortened exposure which supports a higher frame rate for interrogating fast dynamics (Fig. 2), and reduced frame rate which enables longer recording time for investigating long-term variations (Fig. 3).

Furthermore, the temporal resolution of an optical system can be further enhanced without losing spatial resolution through combination with multiplexing techniques [52], and DeepSeMi is readily to mitigate the photodamage due to excessive power dosage. When pushing the frame rate to a limit, a standard device may be capable of imaging ultrafast phenomena like spiking[53] and flagellar locomotion[54] without losing fidelity by using DeepSeMi.

Despite basic exploration has been explored in this manuscript, manifold research can further increase the accessibility of DeepSeMi. By combining with advanced model compression and pruning techniques[55], the computation time of DeepSeMi can be further compressed for high-speed data inference. Training DeepSeMi across a large range of conditions with varied noise and transformations over multiple samples forms a general model, and DeepSeMi in specific conditions with better performance can be swiftly distributed from the basic one with fine-tuning[56].

In short, we believe DeepSeMi provides a robust solution to overcome the shot-noise limitation in fluorescent microscope. Catalyzed by the computational enhancement of DeepSeMi, various organelles and organisms could be safely recorded over a long term in a high spatiotemporal resolution which brings insights into new physiological discoveries.

## Methods

### Network structure

DeepSeMi consisted of six 3D hybrid blind-spot neural networks (four spatial blind spot networks and two temporal blind spot networks) and one feature fusion network (Supplementary Fig. 2). All six hybrid blind-spot networks had the same U-net-like structure for extracting features from input videos. Each hybrid blind-spot network consisted of 14 three 3D convolution layers. The first two layers were 3D eccentric blind-spot convolutional layers with 3 × 3 × 3 sized kernels (Fig. 1c). The encoding path of DeepSeMi was composed by 3D eccentric blind-spot convolutional layers (3×3×3 sized kernels) and MaxPooling layers (2×2×2) alternately. Similarly, the decoding path was implemented by 3D eccentric convolutional layers (3×3×3 sized kernels) and Upsampling layers (2×2×2) alternately. The numbers of input features and output features in each layer were set to 32 to accommodate single-GPU training. The feature fusion network consisted of three 3D convolutional layers with 1×1×1 kernels. The number of input channels of the feature fusion network was 32×6=192 to match the size of concatenated features of the six branch networks, while the number of output channels of the feature fusion network matches the real image and depends on experiments. The loss function of DeepSeMi was a summation of l1 norm and l2 norm, while the learning rate is set to 0.0001.

We usually picked up 1000 patches from noisy videos to form the training set, and the size of each patch was 128×128×32. Good convergence usually could be obtained after 30-50 epochs of training. The entire training process took 10 hours on an NVIDIA 3090 Ti graphics card.

### The concept of eccentric blind-spot convolution and eccentric convolution

The eccentric blind-spot convolution stemming from traditional convolutions plays a significant role in DeepSeMi. Here we illustrate the concept of eccentric blind-spot convolution through derivations. To simplify the description, all following operations are derived in 2D, while the 3D operations can be easily extended.

The traditional discrete convolution (Supplementary Fig. 1a) can be formulated as:

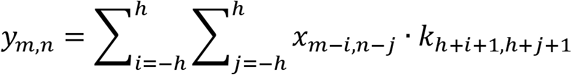

where *y* is the output of the convolution, *x* is the input image, *k* is the kernel of convolution with a size of [2*h* + 1,2*h* + 1]. Note the information of input pixel *x_m,n_* will be transmitted to the output pixel *y_m,n_* in the above traditional convolution process when *i* = 0 and *j* = 0, resulting the noise of input pixel *x_m,n_* will also be kept at the output pixel *y_m,n_*. Training a neural network composed of such convolutional layers in noise-only data will generate trivial results with the identified mapping, and only noisy-clean data pairs or sequential noisy acquisitions can fuel that neural network with the deficiency of self-supervision. To give the neural network the ability to self-supervised denoising, we construct an eccentric blind-spot convolution kernel (Supplementary Fig. 1c), which can be formulated as

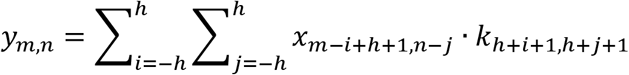

where the symbols are the same as the above equation. With the proposed eccentric blindspot convolution, the noisy information of input pixel *x_m,n_* will not be conserved in the output pixel *y_m,n_*, and information of the output pixel *y_m,n_* can only be estimated from local pixels around the input pixel *x_m,n_*.

In the next we derive the proposed eccentric convolutional filter and explain why it is important to DeepSeMi. In fact, we found that directly combining the aforementioned eccentric blind-spot convolution kernels with traditional convolutional kernels, the blind-spot properties which are the key to ensure the self-supervision would lose. To illustrate that, we concatenate a 2D eccentric blind-spot convolution and a 2D traditional convolution:

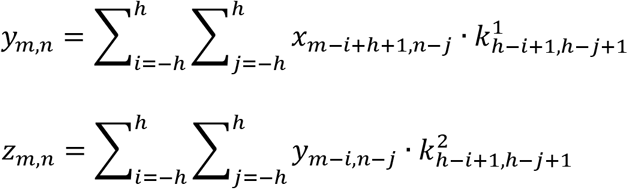

where *x* is the input, *y* is the intermediate variable from the eccentric blind-spot convolutional kernel *k*^1^ and *z* is the output from the traditional convolutional kernel *k*^2^. Both kernels are with size [2*h* + 1,2*h* +1]. It can be easily found that when *h* >0, if

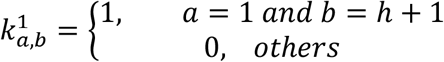

and

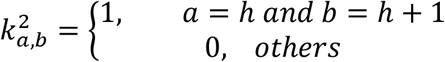

the above formula can be simplified to:

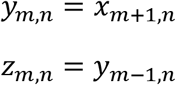

This is equivalent to:

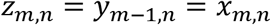

In other words, the original noise pixel *x_m,n_* is directly mapped into an output pixel *z_m,n_* with the same position, indicating the blind-spot properties are dropped. In an extreme condition *h* = 0, such blind-spot properties can be still hold, which explained why we utilized 3D convolutions with kernel size 1 × 1 × 1 in the feature fusion network (FFnet).

To circumvent this shortage, we designed another eccentric convolution which can be formulated as

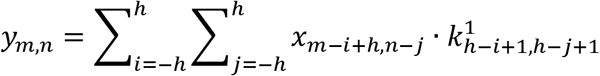

Following the similar derivations as shown above, it can be proved that the blind-spot properties are kept in the combination of fully blind convolutions and eccentric convolutions.

Although the introduction of blind-spot convolutional kernels enabled the neural network to learn denoising without excessive data, the receptive field is limited to only one direction for both the kernels and kernels composited networks. We thus established the hybrid blind-spot network as multiple branches to extract features from different directions, and then fuse these features by feature fusion network (FFnet) to achieve the all-direction-received output result.

#### Time to channel operation

We inserted a time to channel operation [38] at the beginning of the input of the neural network for inputting more temporal information but without obviously increasing the computing time. To achieve that, two times more input frames were input to the network and stacked in the channel dimensions instead of temporal dimensions, which can be quickly squeezed after interacting with the next convolutional kernel. As an example, a video block with a size of C× (T+2F)× H× W was desired to be input, we realigned it to a tensor of size (2FC+C)× T× H× W by multiplexing some frames as the real input of the DeepSeMi.

#### Generation of simulated motion datasets

To fully compare the denoising performance of different algorithms on the video denoising task, we utilized the Moving MNIST dataset as the simulated dataset, which is widely used in the field of computer vision. The images from MNIST handwritten digit database served as the main moving contents in generated videos, while each frame is 256 pixels × 256 pixels in size. In the beginning, we randomly selected 10 handwritten digits to form the basic content, and generated random motions for each of the digits. Then, the whole video was generated frame by frame through keeping shifting the digits in predefined tracks. In order to keep the handwritten digits within the bounds of the video frame, the handwritten digit bounced at the edges of the video frame. The size of the video we usually generate was 256×256×1000.

#### Noise simulation and analysis

We evaluated the performance of DeepSeMi in both Gaussian noise and Poisson noise. The Gaussian noise was simulated by dataset by the *getExperimentNoise* function derived from BM3D [57] with varied noise scales. The Poisson noise was simulated by the *MPG_model* function derived from DeepCAD [33]. We utilized several merits to evaluate the noise scale. Peak signal-to-noise ratio (PSNR) is widely used for measuring the similarity between recovered images and paired ground truth images. The PSNR (in dB) is calculated as:

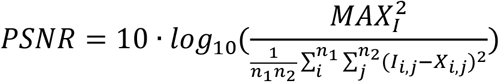

where *X* is a *n*_1_ × *n*_2_ recovered image, *I* is the paired noise-free image. *MAX_I_* is set to 65535 for 16-bit unsigned integer images. Another merit, Signal-to-noise ratio (SNR) is also selected to quantify the image quality after denoising. The SNR (in dB) is calculated as:

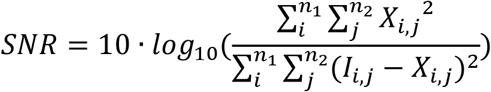

#### Evaluation of photobleaching

Photobleaching represents the inability of the fluorescent protein to emit photons after continuous excitation. To evaluate the photobleaching under different power dosage, we averaged all pixel intensities from the acquired image. To eliminate the influence of the sensor background noise even without fluorescence photons input, we calculated the averaged intensity in a sample-free area, and accordingly updated the averaged intensity across the whole image such that it represents net fluorescence photon flux. We then quantified the speed of photobleaching by fitting the photobleaching curve using an exponential function.

#### Training of organelle segmentation network

As the demand for studying cell biology through microscopic fluorescence imaging increases, it is necessary to utilize automated analysis tools to process massive imaging data in a relatively short time for fertilizing quick experiment iterations. We thereby demonstrated DeepSeMi enhances automated analysis of organelles with high precision and low phototoxicity. We utilized a physics-based machine learning method for organelle segmentation [44]. We simulated both the optical imaging results and segmented ground truth of mitochondria, migrasomes, and retraction fibers based on the morphological characteristics. 1500 paired images were prepared for each organelle. We then built and trained a traditional 2D U-net using the simulated datasets, with the size of the input image of 256×256. It took about 10 minutes on an NVIDIA 3080 Ti graphics card to achieve good convergence results in about 4-10 epochs. The learning rate is set to 0.0001.

We utilized merits of precision, recall, F1-score, and accuracy for segmentation evaluation of the network:

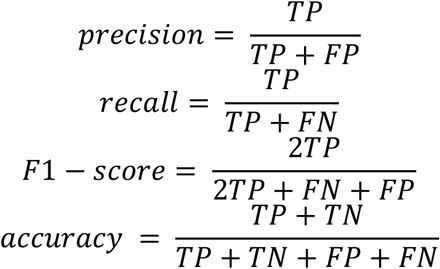

where TP is true positive, TN is true negative, FP is false positive, and FN is false negative.

#### Mitochondria analysis

After mitochondria segmentation through the methods in the above section, the connected regions from the segmented binary masks were detected by *bwlabel* function in MATLAB to accomplish the mitochondrial instance segmentation. The mitochondrial area of each connected region was calculated, and the skeletons and key points of mitochondria were picked up through *bwmorph* function in MATLAB. According to the different topological positions, the key points were classified into junctions or end points. We tracked the mitochondria with Imaris (Oxford Instruments) across recording sessions to indicate the movement state of mitochondria.

#### Cell culture and imaging system

L929 cells and NRK cells were cultured in DMEM (Gibco) medium supplemented with 10% FBS (Biological Industries), 2 mM GlutaMAX and 100 U/ml penicillin-streptomycin in 5% CO2 at 37°C. The PiggyBac Transposon Vector System was used to generate the stably expressing cell line. For L929 cells, Vigofect was used for cell transfection according to the manufacturer’s manual. NRK cells’ transfection was via Amaxa nucleofection using solution T and program X-001. 35 mm confocal dishes were precoated with fibronectin (10 mg/ml) at 37°C for 1 hour. Cells were cultured in fibronectin-precoated confocal dishes for 4 hours before imaging.AX2 axenic strain cells were provided by the Jeffrey G Williams laboratory (Cell and Developmental Biology, College of Life Sciences, University of Dundee, UK). The AX2 WT cells and the derived cell line were cultured in HL5 medium (Formedium # HLF2), supplemented with antibiotics, at 22°C. The plasmids pDM323 and pDM451 were provided by the Huaqing Cai laboratory (National Laboratory of Biomacromolecules, Institure of Biophysics, Chinese Academy of Sciences, China). The DNA fragments encoding dajumin and cAR1 were PCR-amplified and cloned into the overexpressing plasmids.

*C. elegans* stably overexpressing OSM-3-GFP were provided by the Guangshuo Ou laboratory (School of Life Sciences, Tsinghua University, China). We cultivated *C. elegans* on nematode growth medium agar plates seeded with the *Escherichia coli* OP50 at 20 °C. For live cell imaging, worms were anesthetized with 1 mg/mL levamisole and mounted on 3% agarose pads at 20 °C.

The *Tg(mpeg1.1:PLMT-eGFP-caax)* transgenic zebrafish was provided by Boqi Liu. All adult zebrafish were kept in a water-circulating system at 28.5 °C. Fertilized eggs were raised at 28.5 °C in Holtfreter’s solution. The embryos were embedded in 1% low-melting-point agarose for live-cell imaging. The use of all zebrafish adults and embryos was conducted according to the guidelines from the Animal Care and Use Committee of Tsinghua University.

All imaging experiments in this research were based on a Nikon A1 confocal microscope (Bioimaging center, School of Life Sciences, Tsinghua University, China). All cellular imaging was conducted by a 100 × objective (NA 1.45, oil immersion). A 10x objective (10×, NA 0.45, air) was used to capture the global image of *C. elegans* and zebrafish. Two-photon imaging was conducted with a customized two-photon imaging system under a commercial objective (25×, NA 1.05, XLPLN25XWMP2, Olympus).

## Data availability

All relevant data that support the findings of this study are available from the corresponding authors upon reasonable request.

## Code availability

Our DeepSeMi can be found at https://github.com/GuoxunZhang-PhD/DeepSeMi.

## Acknowledgments

We would like to acknowledge the assistance of SLSTU-Nikon Biological Imaging Center for the assistance of using Nikon A1 confocal microscopy. We thank the assistance of the Imaging Core Facility, Technology Center for Protein Sciences, Tsinghua University for the assistance of using Imaris for organelle tracking. We thank Hao Zhu for providing us with *C. elegans.* We thank Haifeng Jiao for providing us with several stable cell lines and suggestions. This work was supported by NSFC (No. 62088102, 62222508 and 62071272) and MOST(No.2020AA0105500).

## Author contributions

Q.D., L.Y. and J.W. conceived the DeepSeMi project and revised the manuscript. G.Z. implemented the DeepSeMi pipeline, performed simulations and analyzed the imaging results. X.L. cultivated biological cells, conducted confocal imaging experiments and revised the manuscript. Y.Z. provided constructive suggestions and wrote the manuscript. X.H. and X.L. revised the manuscript. J.Y. provided *Dictyostelium* cells and B.L. provided zebrafish.

