## Supplementary Materials for "Bio-friendly long-term subcellular dynamic recording by self-supervised image enhancement microscopy"

| | |
|---|---|
| **Supplementary Figure 1** | Comparison of the proposed blind-spot convolutional filter with the traditional 2D convolution. |
| **Supplementary Figure 2** | The network structure of DeepSeMi. |
| **Supplementary Figure 3** | The denoising benchmark of DeepSeMi and other methods on Gaussian-noise corrupted Moving MNIST datasets over different noise scales. |
| **Supplementary Figure 4** | The denoising benchmark of DeepSeMi and other methods on Gaussian-noise corrupted Moving MNIST datasets over different content speeds. |
| **Supplementary Figure 5** | Comparison of DeepSeMi with DeepCAD and DeepInterpolation on Gaussian-noise corrupted Moving MNIST datasets over different content speeds. |
| **Supplementary Figure 6** | DeepSeMi corrects motion artifacts that defile DeepCAD. |
| **Supplementary Figure 7** | The denoising benchmark of DeepSeMi and other methods on Poissionian-noise corrupted Moving MNIST datasets. |
| **Supplementary Figure 8** | Evaluation of generalization ability of DeepSeMi on simulated datasets. |
| **Supplementary Figure 9** | Mitochondria membrane-trained DeepSeMi effectively works on mitochondria matrix and cell membrane imaging. |

| | |
|---|---|
| **Supplementary Figure 10** | Denoising of Tom20-GFP-mCherry-labeled mitochondrial via DeepSeMi. |
| **Supplementary Figure 11** | DeepSeMi effectively enhances SNR of triple-color labeled multi organelles. |
| **Supplementary Figure 12** | Laser power calibration on the Nikon A1 confocal microscopy. |
| **Supplementary Figure 13** | Evaluation of photobleaching of mitochondria under different laser dosages. |
| **Supplementary Figure 14** | DeepSeMi helps automated segmentation and skeletonization of mitochondria under low power dosage. |
| **Supplementary Figure 15** | DeepSeMi enables high-SNR imaging of tri-color labeled L929 cells in low light. |
| **Supplementary Figure 16** | 15-fold increment of photon budgets by DeepSeMi. |
| **Supplementary Figure 17** | DeepSeMi significantly enhances organelle imaging results with the dye dilution. |
| **Supplementary Figure 18** | Significant photobleaching brought by dual-color confocal imaging. |
| **Supplementary Figure 19** | Statistics of mitochondrial segmentation and skeletonization under different illumination powers with and without DeepSeMi enhancement. |

| | |
|---|---|
| **Supplementary Figure 20** | Automated analysis of recorded mitochondria with DeepSeMi enhancement. |
| **Supplementary Figure 21** | DeepSeMi-enhanced imaging results of L929 cells treated with Lat-A. |
| **Supplementary Figure 22** | DeepSeMi unveiled migrating cells interacting with a migrasome, producing migrasomes, and expelling mitochondria in low light dosage. |
| **Supplementary Figure 23** | Evaluation of phototoxicity in imaging *Dictyostelium* cells. |
| **Supplementary Figure 24** | Evaluation of phototoxicity in imaging Dictyostelium cells with a bright-field microscope imaging. |
| **Supplementary Figure 25** | DeepSeMi enables high-SNR imaging of contractile vacuole generation in photosensitive *Dictyostelium* cells. |
| **Supplementary Figure 26** | DeepSeMi enhanced cellular observation in scattering *C. elegan* in vivo. |
| **Supplementary Figure 27** | DeepSeMi enhances observation of zebrafish larvae in a low light dosage. |
| **Supplementary Figure 28** | DeepSeMi effectively recovers functional data on open-sourced two-photon Neurofinder datasets. |
| **Supplementary Figure 29** | Evaluation of DeepSeMi on hybrid high and low-SNR functional imaging. |

| | |
|---|---|
| **Supplementary Figure 30** | Evaluation of DeepSeMi on hybrid high and low-SNR dendritic imaging. |

**Video captions**

| | |
|---|---|
| **Supplementary Video 1** | Evaluation and segmentation of DeepSeMi enhancement over a triple-color labeled L929 cell in low light. First 18 seconds: comparison of the raw noisy captured video (top) and the DeepSeMi-enhanced video (bottom) of a triple-color labeled L929 in a commercial confocal microscope, with a panel indicating intensity profiles along the green line attached in the bottom right corner.18 to 37 seconds: comparison of raw (top) and DeepSeMi-enhanced zoom-in video outlined by the white box. 38 seconds to the end: comparison of mitochondria segmentation and keypoint detection of raw (top) and DeepSeMi-enhanced (bottom) video (Methods), where red points for junction points, yellow points for endpoints, green lines for mitochondrial skeletons, and gray area for mitochondrial segments. |
| **Supplementary Video 2** | Evaluation of photobleaching of mitochondria under different laser dosages. Mitochondria in L929 cells were captured by a commercial confocal microscopy with five different excitation laser intensity (0.5% 14.6 μW, 1% 23.1 μW,2% 45.3 μW, 4% 80.4 μW, 8% 152.3 μW). The first row presented the raw (top) and DeepSeMi-enhanced (bottom) mitochondria movie. The second row presented corresponding photobleaching curves of each power dosage (Methods). |

| | |
|---|---|
| **Supplementary Video 3** | Evaluation of DeepSeMi enhancement in a quadruple-color labeled L929 cell in low light over 13,000 frames. The L929 cell was imaged in a commercial confocal microscope for a half-hour long session and presented in the left, and intensity profiles along the green and yellow lines were dynamically presented on the right, where the first and third rows for raw and the second and fourth rows for DeepSeMi. The first 12 seconds: raw noisy captured video of a L929 cell. 12 seconds to the end: DeepSeMi enhanced video of a L929 cell. Frame numbers and time stamps were annotated on the right bottom. |
| **Supplementary Video 4** | Evaluation of DeepSeMi enhancement in observation of cell migrations in low light over 12 hours. Two L929 cells were imaged in a commercial confocal microscope over 80,000 frames. The first 40 seconds: raw (left) and DeepSeMi-enhanced (right) videos representing migrations of two L929 cells in a global view. 40 seconds to the end: raw (left) and DeepSeMi-enhanced (right) video representing migrations of two L929 cells in a zoom-in view, where generation of migrasomes was clearly presented. Frame numbers and time stamps were annotated on the right bottom. |
| **Supplementary Video 5** | Evaluation of DeepSeMi enhancement in observation of retractosomes generation. Two L929 cells were imaged in a commercial confocal microscope for over 24,000 |

| | |
|---|---|
| | seconds. Raw (left) and DeepSeMi-enhanced (right) videos representing generation of retractosomes were presented. Frame numbers and time stamps were annotated on the right bottom. |
| **Supplementary Video 6** | Evaluation of DeepSeMi enhancement in observation of intercell interactions in low light over 2 hours. Two L929 cells were imaged in a commercial confocal microscope. The first 40 seconds: raw (top) and DeepSeMi-enhanced (bottom) video representing the generation of migrasomes on retraction fibers in a global view. 40 seconds to 1 minute 5 seconds: raw (top) and DeepSeMi-enhanced (bottom) videos representing the generation of migrasomes on retraction fibers in a zoom-in view. 1 minute 5 seconds to 1 minute 20 seconds: raw (top) and DeepSeMi-enhanced (bottom) video representing interactions between a cell and a migrasome. 1 minute 20 seconds to the end: raw (top) and DeepSeMi-enhanced (bottom) videos representing a long-distance movement of mitochondria. Frame numbers and time stamps were annotated on the right bottom. |
| **Supplementary Video 7** | Evaluation of DeepSeMi enhancement in observation of *Dictyostelium* cells in low light. Three *Dictyostelium* cells were imaged in a commercial confocal microscope over 1800 seconds and presented in 0~192 seconds, 0~276 seconds, and 7~1308 seconds, respectively. |

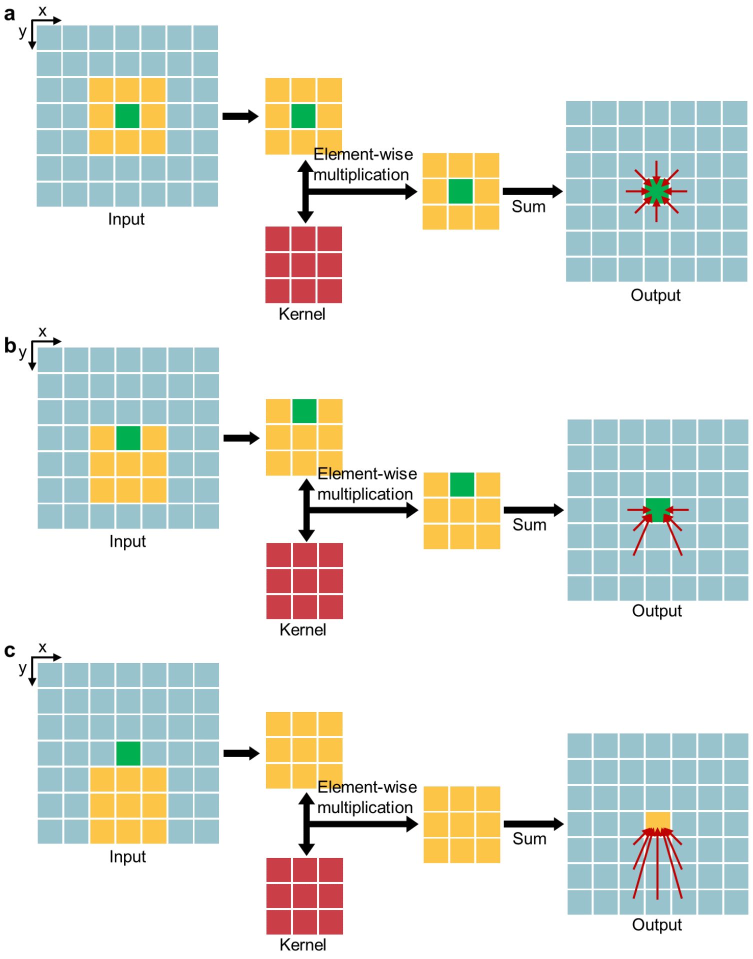


**Supplementary Figure 1. Comparison of the proposed eccentric convolutional filter with the traditional 2D convolution**. **a**, Processing of the traditional convolutional filter. The convolution procedure takes place around the target pixel (green) with a kernel size of 3x3 (red), and the receptive field is marked by yellow pixels and the green pixel. **b**, Processing of the eccentric convolutional filter. The symbols are the same as **a**. Note the receptive field is biased off the target pixel but does not entirely miss the target pixel. Due to the concatenated fashion of the convolutional neural network, the receptive field of following convolutional layers is also limited. **c**, Processing of the eccentric blind-spot convolutional filter. The symbols are the same as **a**. Note the receptive field is entirely off the target pixel.

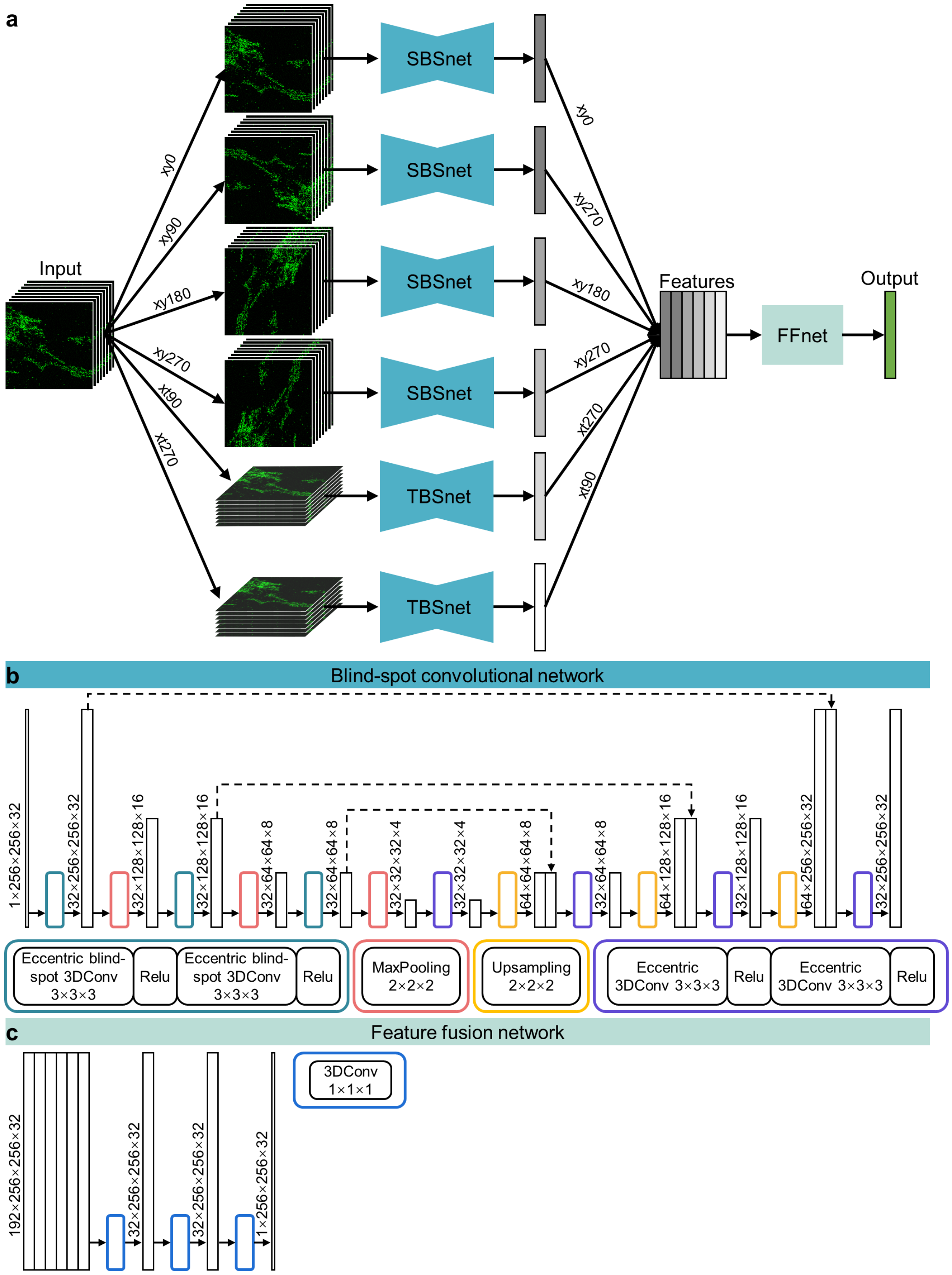


**Supplementary Figure 2. The network structure of DeepSeMi**. **a**, Six branches of hybrid blind-spot 3D neural network which has different preferential directions of reception field compose the DeepSeMi. Among the six branches, four networks are spatial blind-spot 3D neural networks (SBSnets) and two networks are temporal blind-spot 3D neural networks (TBSnets). The four SBSnets and the two TBSnets share the same parameters, respectively. Output features from the six branches are concatenated and input to a feature fusion network (FFnet) for the final output. **b**, Detailed structure of the blind-spot 3D neural network in each branch. **c**, Detailed structure of the feature fusion network. Feature fusion network takes advantage of 1×1×1 3D convolutions to merge features from each branch.

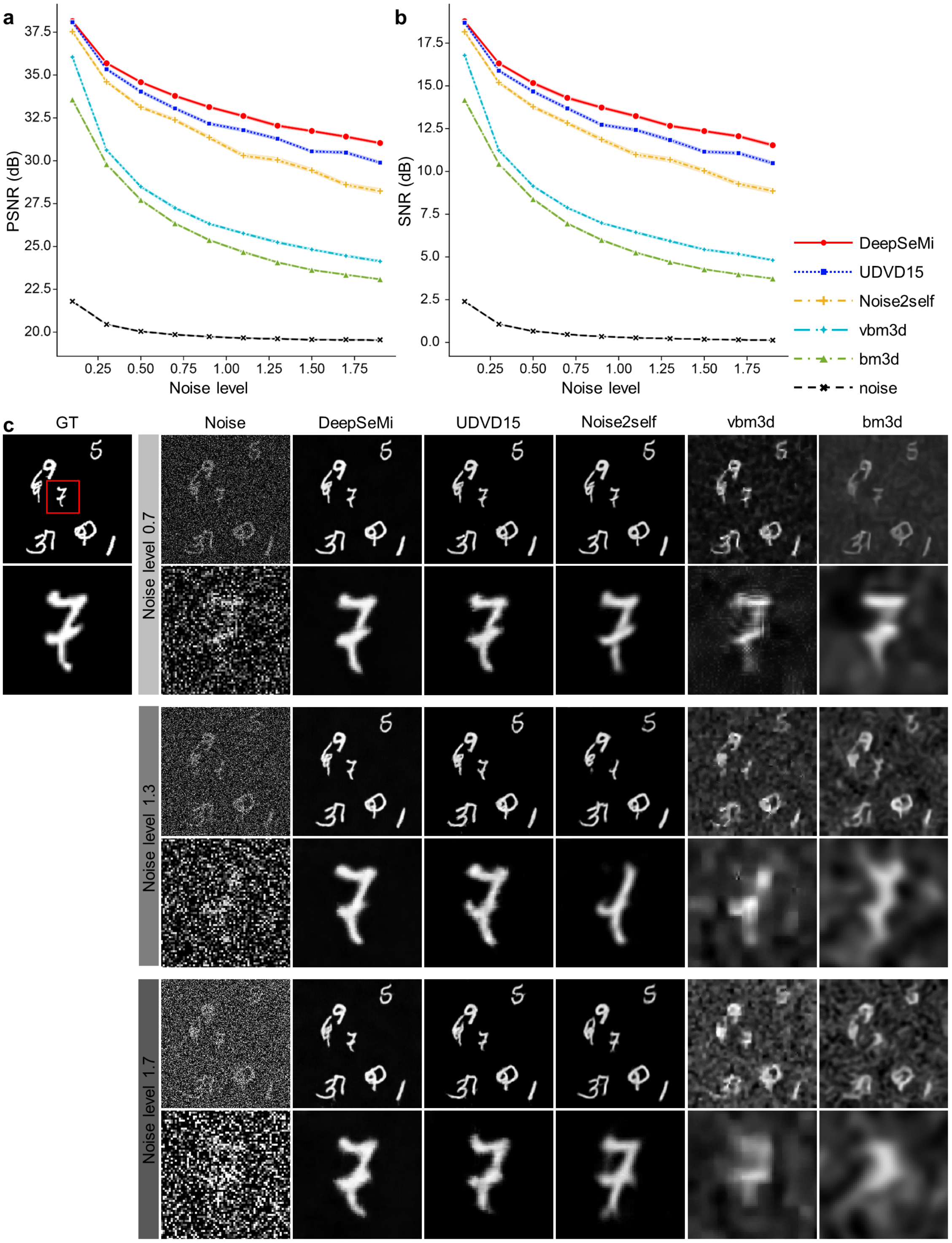


42 **Supplementary Figure 3. The denoising benchmark of DeepSeMi and other methods**
43 **on Gaussian-noise corrupted Moving MNIST datasets over different noise scales. a-**
44 **b,** Peak-signal-to-noise-ratio (PSNR) and signal-to-noise-ratio (SNR) comparisons of
45 DeepSeMi, UDVD15 [1], Noise2Self [2], VBM3D [3], and BM3D [4] at different noise

levels. The motion speed is set as 5, which means the handwritten digits in the next frame are shifted by 5 pixels relative to the previous frame. **c**, Exemplary denoising results of DeepSeMi and other methods over three noise levels. The first row on each noise level represents the full field and the second row represents the zoom-in area marked by the red dashed box (in this case, the number “7”).

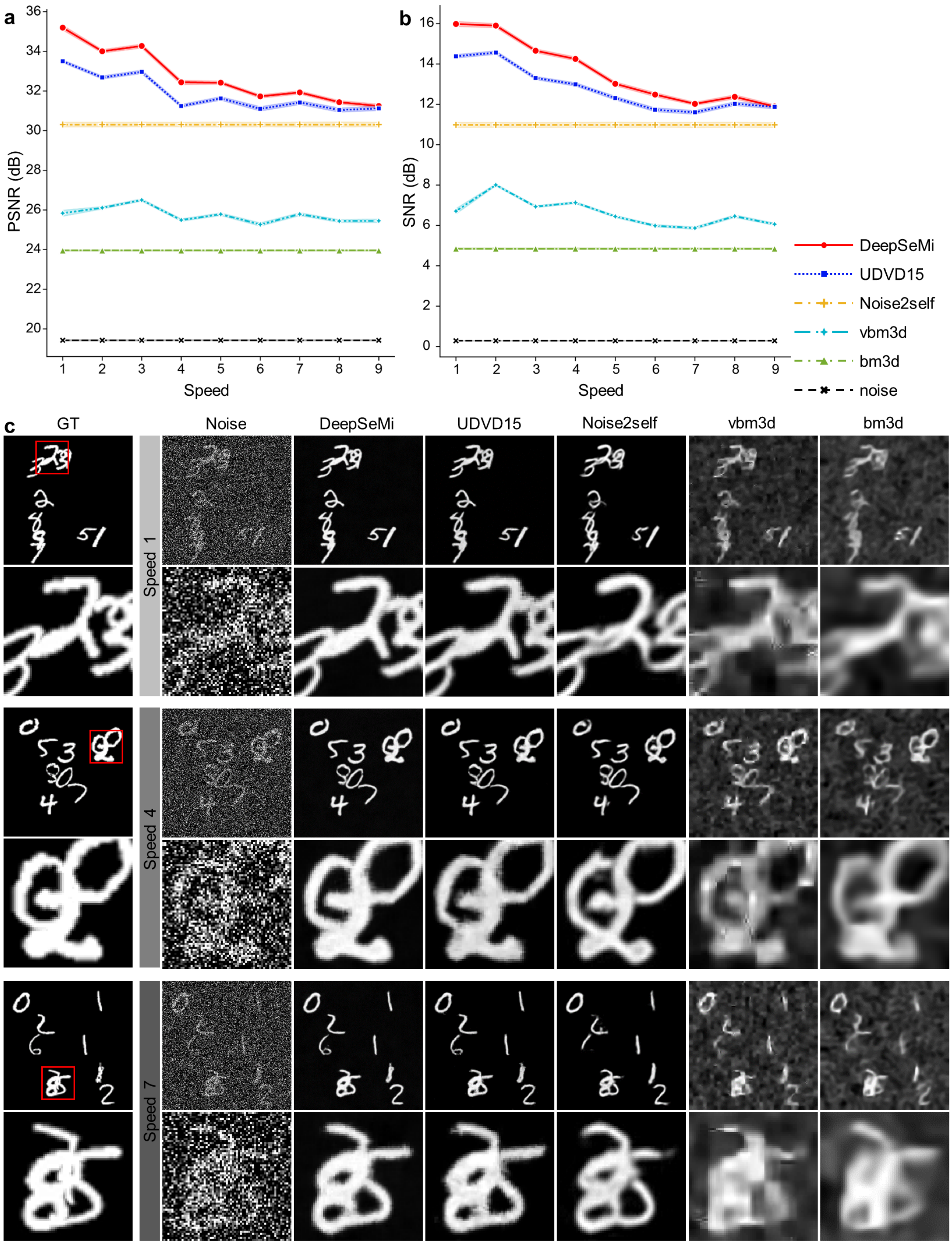


**Supplementary Figure 4. The denoising benchmark of DeepSeMi and other methods on Gaussian-noise corrupted Moving MNIST datasets over different content speeds.** **a-b**, PSNR and SNR comparisons of DeepSeMi, UDVD15 [1], Noise2Self [2], VBM3D [3], and BM3D [4] at different content motion speeds. The motion speed N is defined as the relative shift step in pixel between adjacent frames of each handwritten digit. **c**,

Exemplary denoising results of DeepSeMi and other methods over three motion speeds. The first row on each speed represents the full field and the second row represents the zoom-in area marked by the red dashed box.

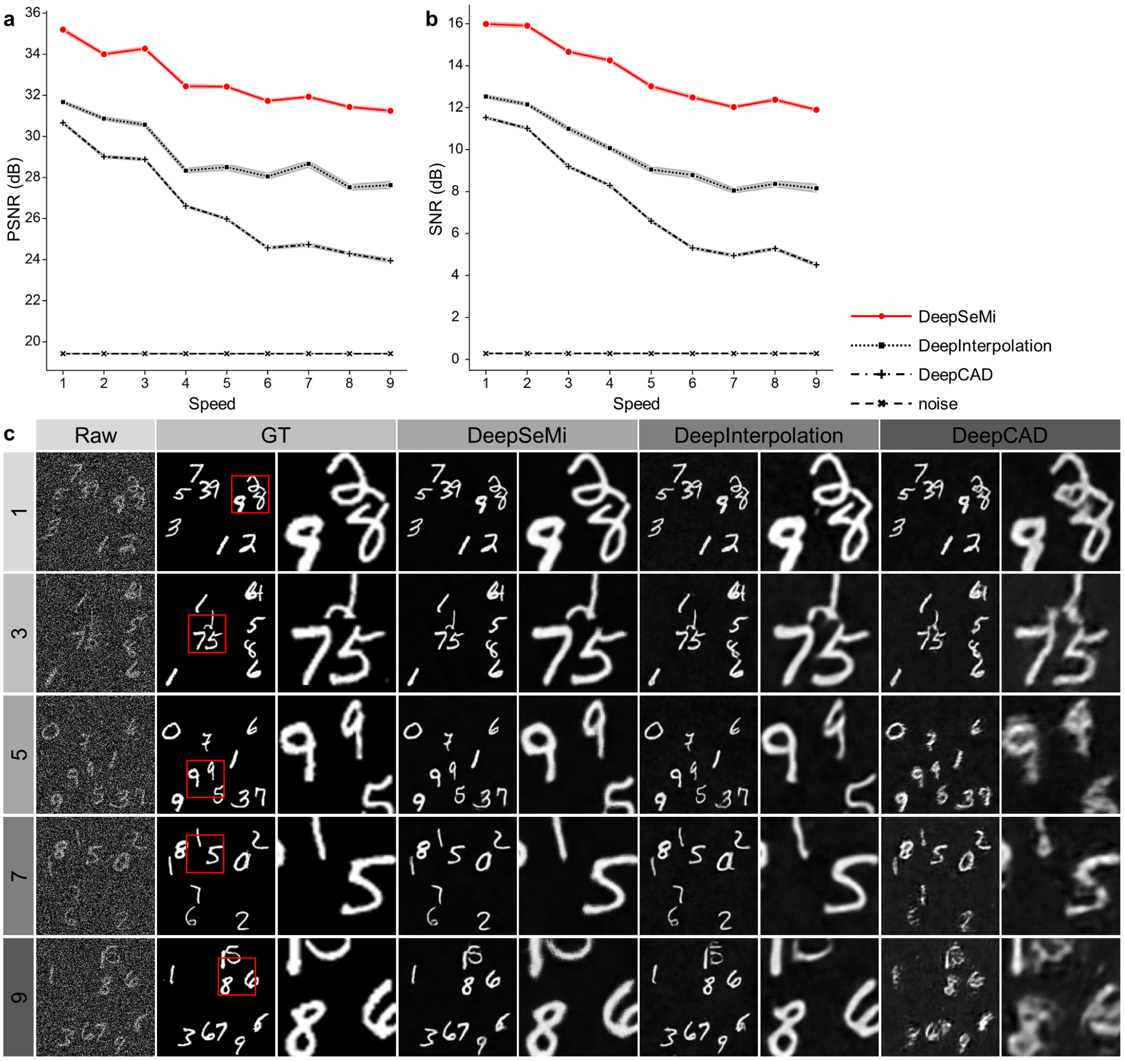


**Supplementary Figure 5. Comparison of DeepSeMi with DeepCAD and DeepInterpolation on Gaussian-noise corrupted Moving MNIST datasets over different content speeds**. **a-b**, PSNR and SNR comparisons of DeepSeMi, DeepCAD [5], and DeepInterpolation [6]. The motion speed N is defined as the relative shift step in pixel between adjacent frames of each handwritten digit. The noise level is set as 1.1. **c**, Exemplary denoising results of DeepSeMi and other methods over three motion speeds. The first row on each speed represents the full field and the second row represents the zoom-in area marked by the red dashed box.

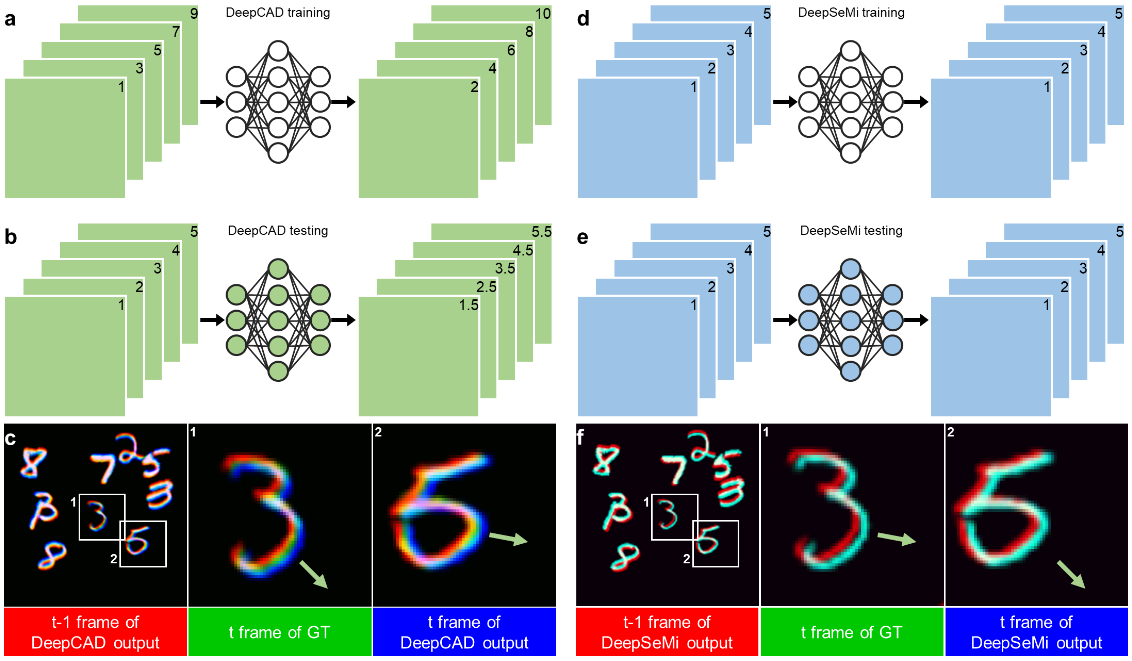


**Supplementary Figure 6. DeepSeMi corrects motion artifacts that defile DeepCAD.** **a**, Schematic diagram represents the DeepCAD training process with the frame number labeled. **b,** Schematic diagram represents the DeepCAD testing process. Note the inferred frames are the interpolations between captured adjacent frames that are not physically captured. **c,** Motion artifacts of DeepCAD on Moving MNIST datasets. A DeepCAD model was trained and tested on Moving MNIST datasets (left), and temporal-color coded digits were presented in zoom-in panels (right). The motion directions are labeled by the arrows. The red channel represents the output of DeepCAD in frame *t-1*, the green channel represents the ground truth in frame *t*, and the blue channel represents the ground truth in frame *t* as a reference. Blue and green channels were obviously separated, indicating prediction bias in DeepCAD over moving contents. **d-f**, The same as **a-c** but from results by DeepSeMi. Blue and green channels were closely matched in **f**, indicating no motion artifacts generated by DeepSeMi.

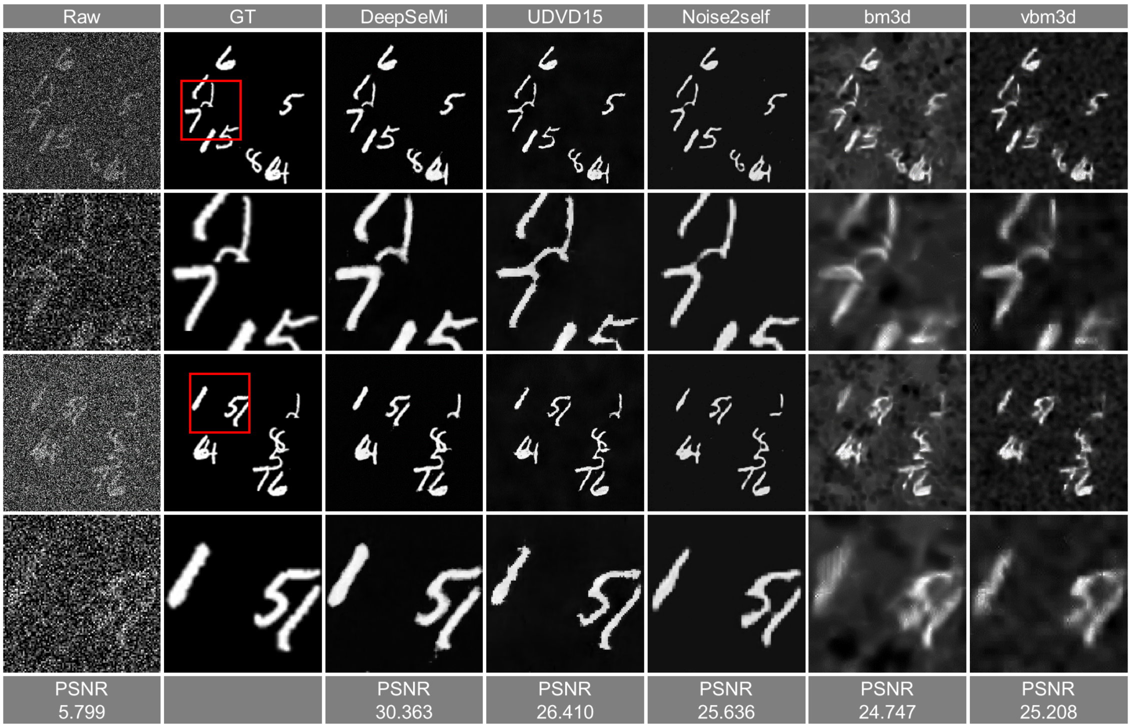


**Supplementary Figure 7. The denoising benchmark of DeepSeMi and other methods on Poission-noise corrupted Moving MNIST datasets.** We test the denoising performance of DeepSeMi, UDVD 15 [1], Noise2self [2], BM3D [4], and VBM3D [3]. The motion speed is set as 5. The second and fourth rows are magnified images of the red box in the first and third rows, respectively. Among these six algorithms, DeepSeMi has the best denoising performance with clean details.

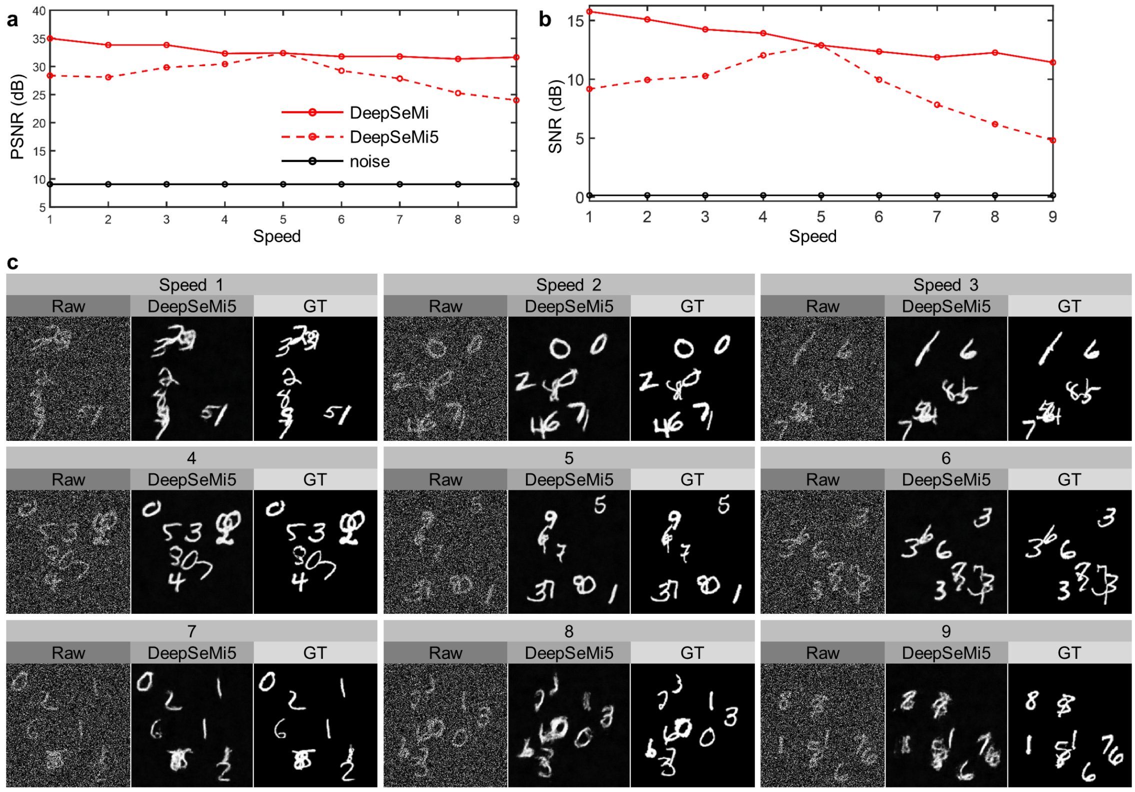


**Supplementary Figure 8. Evaluation of generalization ability of DeepSeMi on simulated datasets**. DeepSeMi is trained through the simulation data with a speed of 5 (the handwritten digit in the next frame is shifted by 5 pixels relative to the previous frame) and a noise level of 1.1, and is termed as DeepSeMi5. The trained DeepSeMi5 is used to denoise simulation data with different motion speeds in the following panels. **a-b**, The generalization ability of DeepSeMi5 over different content speeds but the same noise level measured by PSNR and SNR comparison. The red solid line represents results by DeepSeMi trained on datasets with current content speed and noise level. The red dashed line represents results by DeepSeMi5. The black solid line represents the characteristics of noise. **c**. Exemplary plots of denoising results of DeepSeMi5 over content speed 1 to 9.

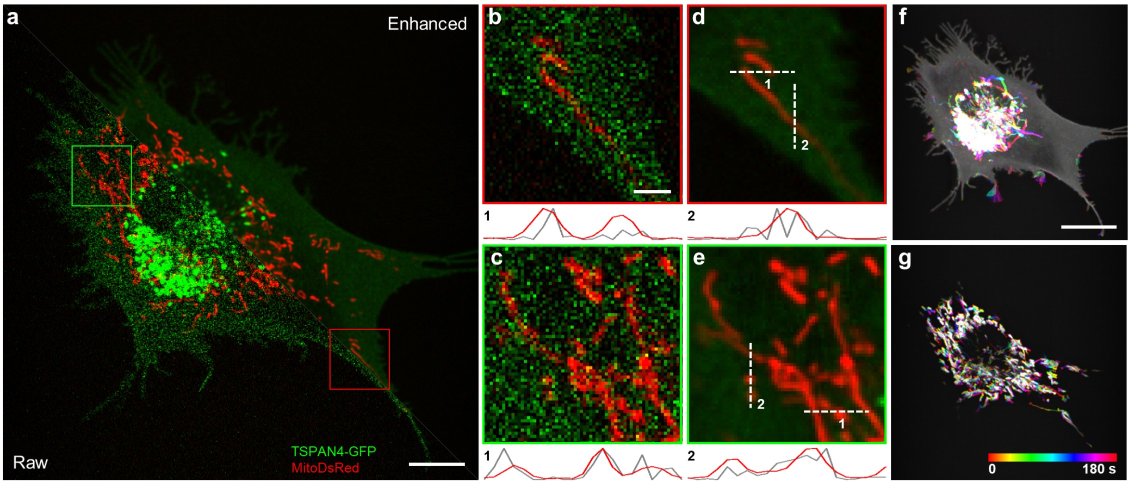


**Supplementary Figure 9. Mitochondrial membrane-trained DeepSeMi effectively works on mitochondria matrix and cell membrane imaging. a**, Raw (left) and DeepSeMi enhanced (right) images of simultaneously captured cell membrane (green) and mitochondria matrix (red). We trained the DeepSeMi only on the experimental data of mitochondrial membrane but tested it on both cell membrane and mitochondria matrix. Scale bar, 10 µm. **b-e**, Zoom-in panels of the box-enclosed regions in **a**. Intensity profiles along dashed lines are plotted at the bottom. Scale bar, 2 µm. **f-g**, Temporal-color coded cell membrane and mitochondrial dynamics, respectively. Scale bar, 20 µm.

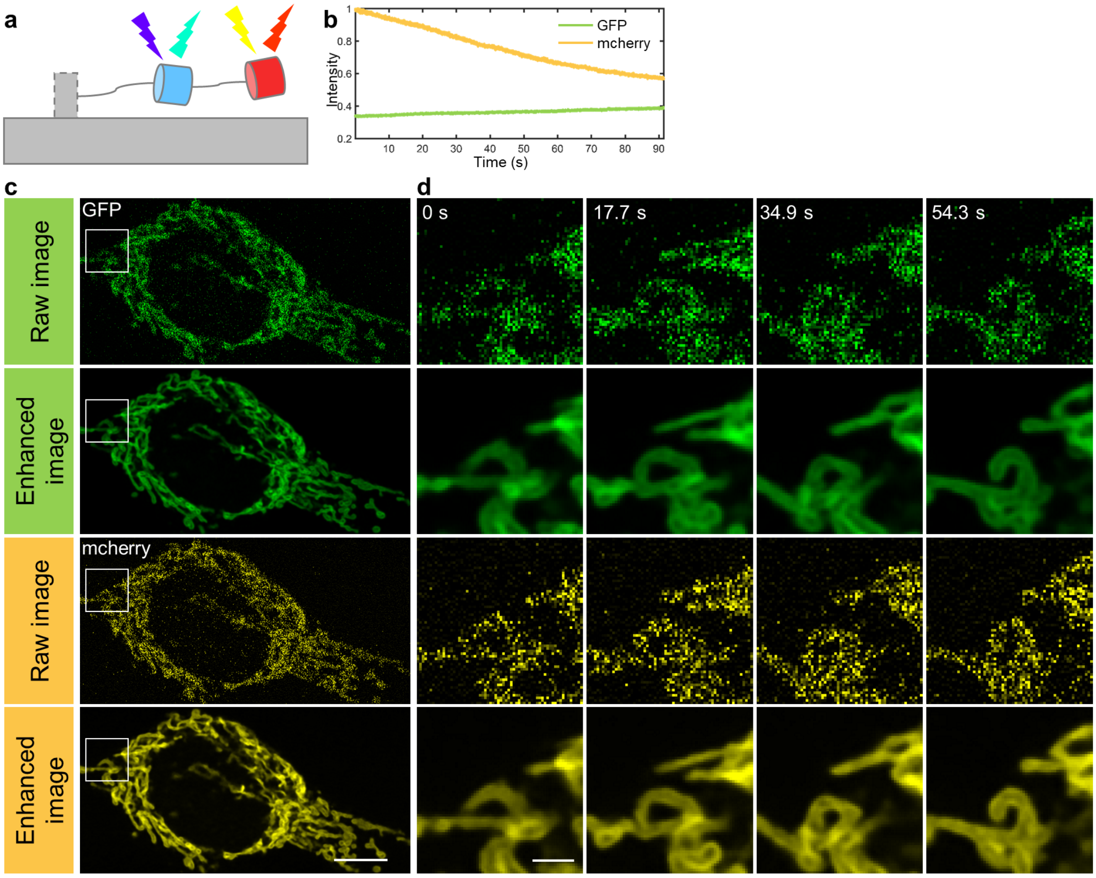


**Supplementary Figure 10. Denoising of Tom20-GFP-mCherry-labeled mitochondria via DeepSeMi**. **a**, Schematic diagram of Tom20-GFP-mCherry-labeled mitochondria which carries two fluorescence tag. **b**, Different photobleaching rates of GFP and mCherry in the co-labeled mitochondria. **c**, Raw and DeepSeMi denoised results labeled by GFP and mCherry. Scale bar, 10 µm. **d**, Zoom-in view of the white box in **c** at different time points. Scale bar, 2 µm.

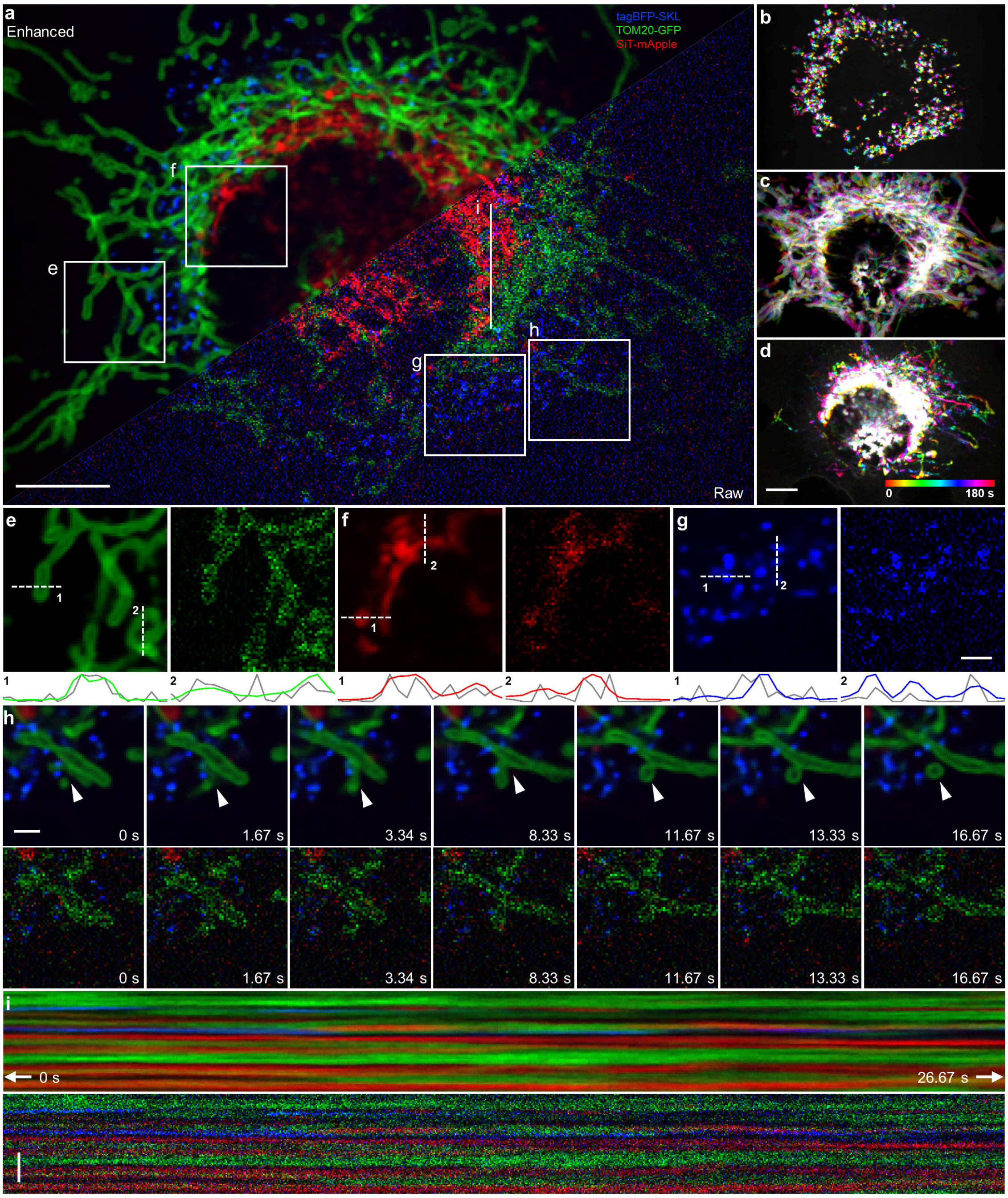


**Supplementary Figure 11. DeepSeMi effectively enhances SNR of triple-color labeled multiple organelles. a**, Raw (right) and DeepSeMi denoised (left) images of mitochondria (green), peroxisomes (blue), and Golgi (red) in an L929 cell in 1,800 frames per channel during 180 seconds. Scale bar, 10 µm. **b-d**, Temporal color coding of denoised mitochondria, peroxisomes, Golgi images to reflect organelle dynamics. Scale bar, 10 µm. **e-g**, Zoom-in panels of white boxes marked in **a**. The membranous structures of mitochondria and the punctate structures of peroxisome were clearly retrieved by DeepSeMi, and the structure profiles along the white dashed lines (colorful, bottom of the figure) are more reasonable compared to raw images (dark, bottom of the figure). Scale bar, 2 μm. **h**, Time-lapse presentation of a vesicle fission event by DeepSeMi (top) and raw (bottom), where a globular mitochondria split from another rod-shaped mitochondrion. Scale bar, 2 μm. **i**, Kymographs (y-t view) of raw images (bottom) and DeepSeMi enhanced images (top) along the white solid line in **a**. Scale bar, 4 μm.

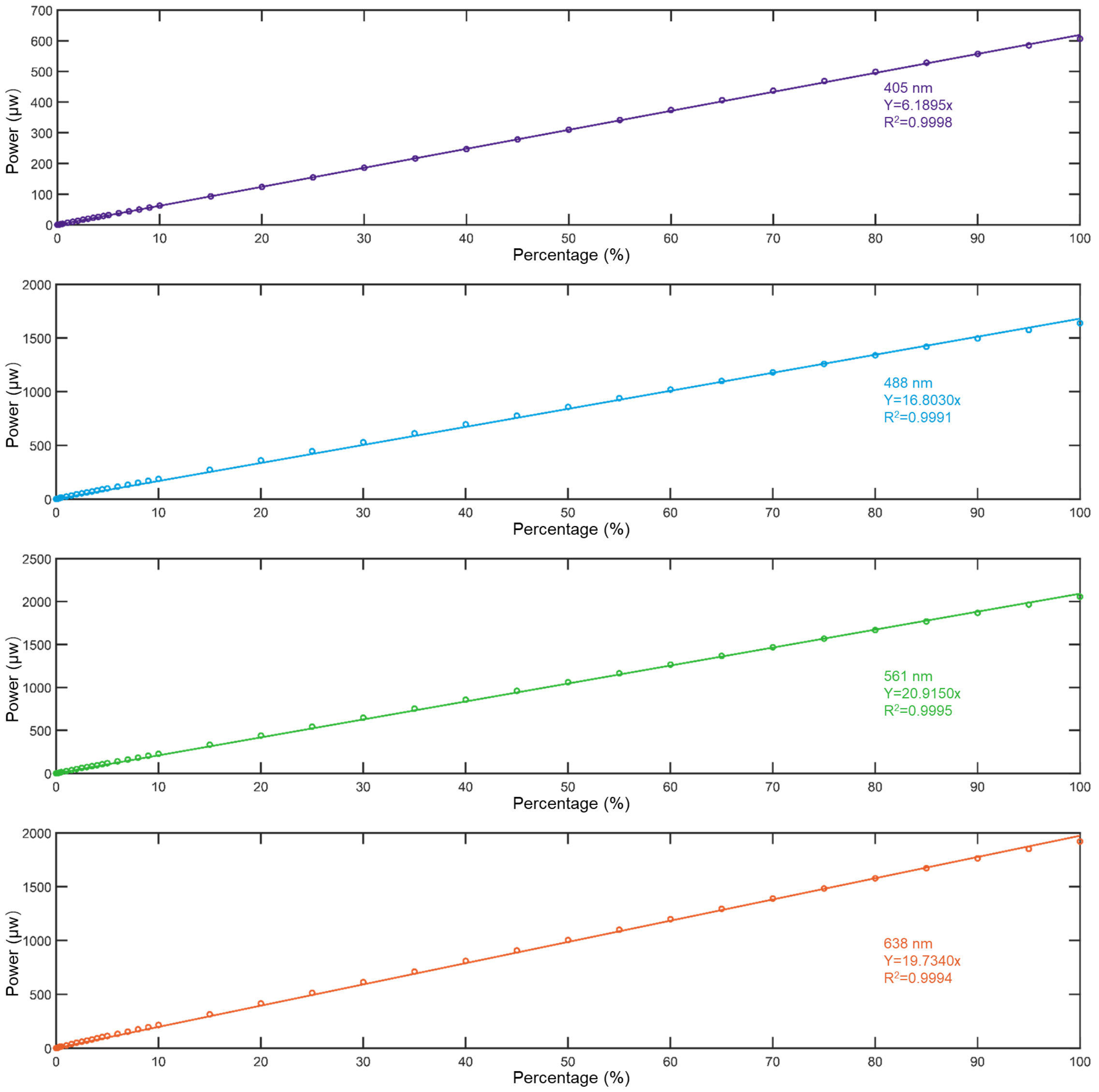


**Supplementary Figure 12. Laser power calibration on the Nikon A1 confocal microscopy**. We calibrated the laser power of the commercial confocal microscope (Nikon A1) which was used for experiments presented in this research. The laser power was measured at the exit of the objective through a power meter (Thorlabs, PM100D) for four wavelengths separately (405 nm, 488 nm, 561 nm, 638 nm). The linearity of powers across all wavelengths was higher than 0.999.

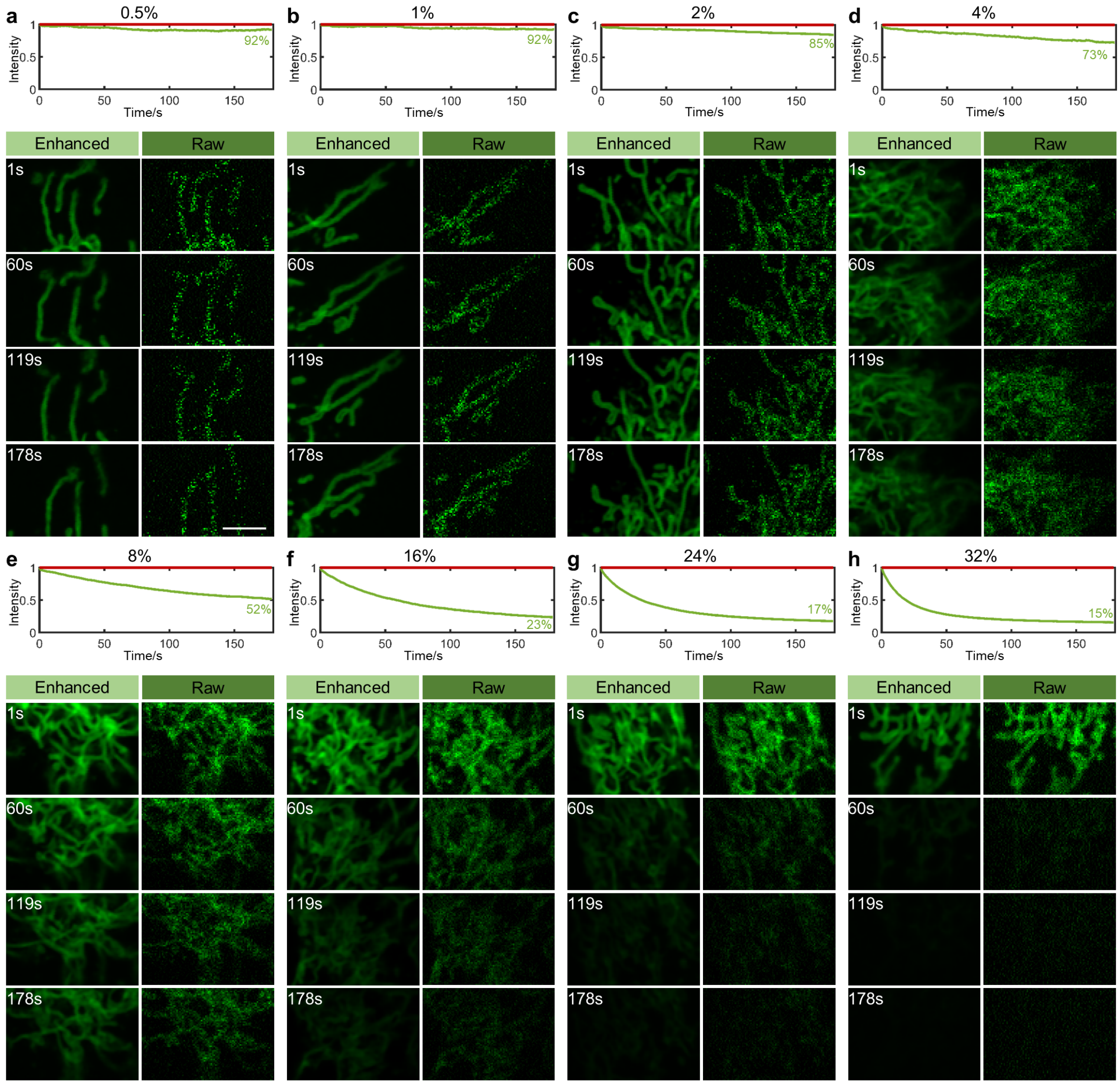


**Supplementary Figure 13. Evaluation of photobleaching of mitochondria under different laser dosages.** All images are captured under the resonant-scanning mode by commercial confocal microscopy (Nikon A1). **a-h**, Intensity statistics and time-lapse images of mitochondria under different laser intensities (488 nm: 0.5%, 1%, 2%, 4%, 8%, 16%, 24%, 32%). For each laser intensity, 5,400 frames are captured within a 3-minute session window. Top, intensity fluctuations during the laser illumination, where the red line is the reference representing zero photobleaching. Bottom, DeepSeMi enhanced (left) and raw (right) data at four different time points. Scale bar, 5 µm.

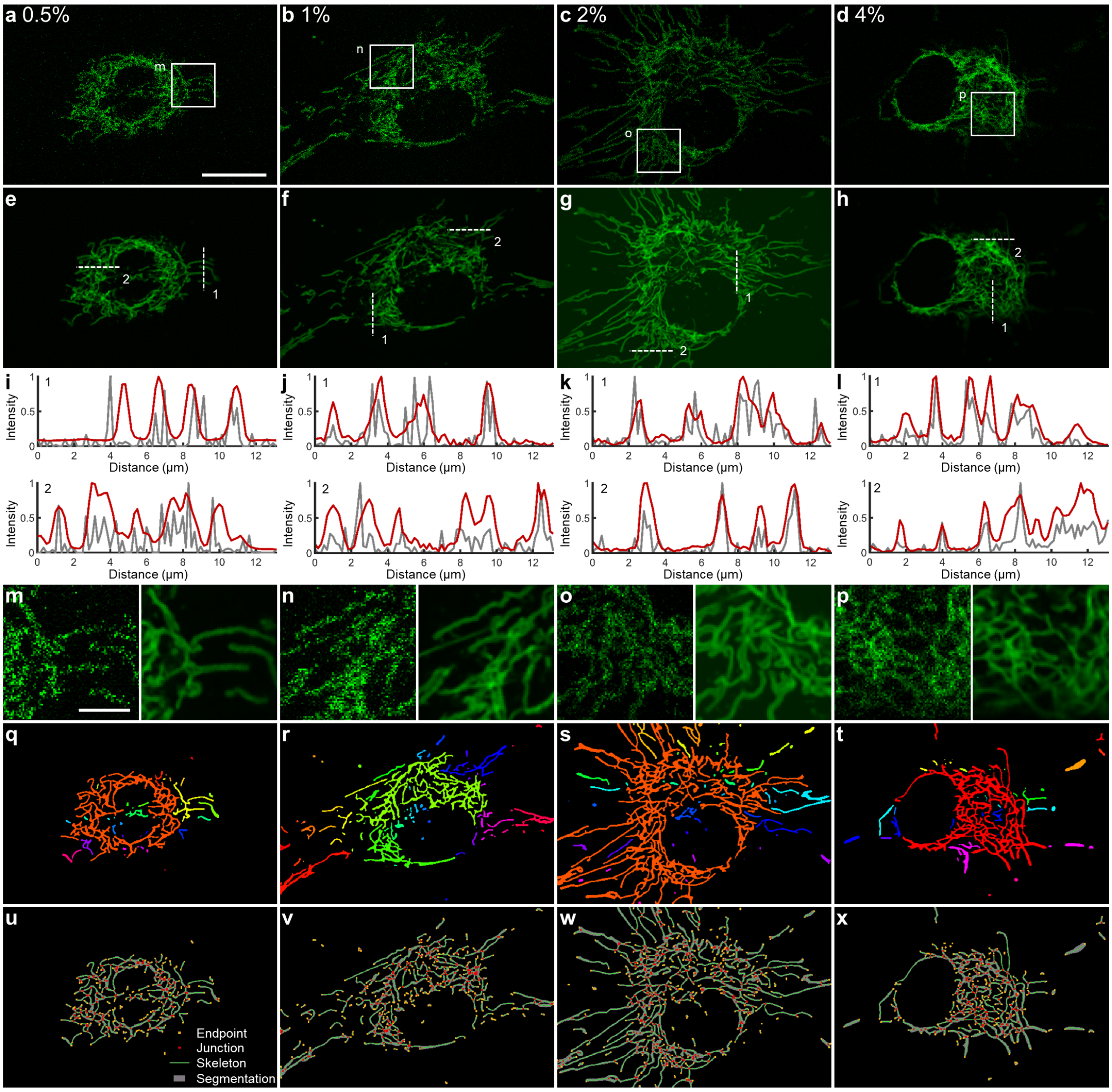


**Supplementary Figure 14. DeepSeMi helps automated segmentation and skeletonization of mitochondria under low power dosage. a-d**, Raw captures of mitochondria at four light intensities (488 nm:0.5%, 1%, 2%, 4%). Scale bar, 20 μm. **e-h**, DeepSeMi enhanced results of mitochondrial corresponding to **a-d**, respectively. It is obvious that DeepSeMi results in clear structures and cleaner backgrounds across all intensities with delicate mitochondrial details recovered. **i-l**, Intensity profiles along the white dashed line in **e-h**, respectively. The profiles by DeepSeMi enhancement (red) are more reasonable. **m-p**, Zoom-in panels of white boxes in **a-d**, respectively. DeepSeMi reunites fragmented structures due to noise contaminations and unveils rich structures of mitochondria. Scale bar, 5 μm. **q-t**, Instance segmentation of DeepSeMi enhanced mitochondrial images through a simulation-supervision machine learning algorithm (Methods). Different colors represent different connected regions. **u-x**, Segmentation (gray), skeletonization (green), and key point detection (yellow for end point, red for junction point) after mitochondrial segmentation (Methods).

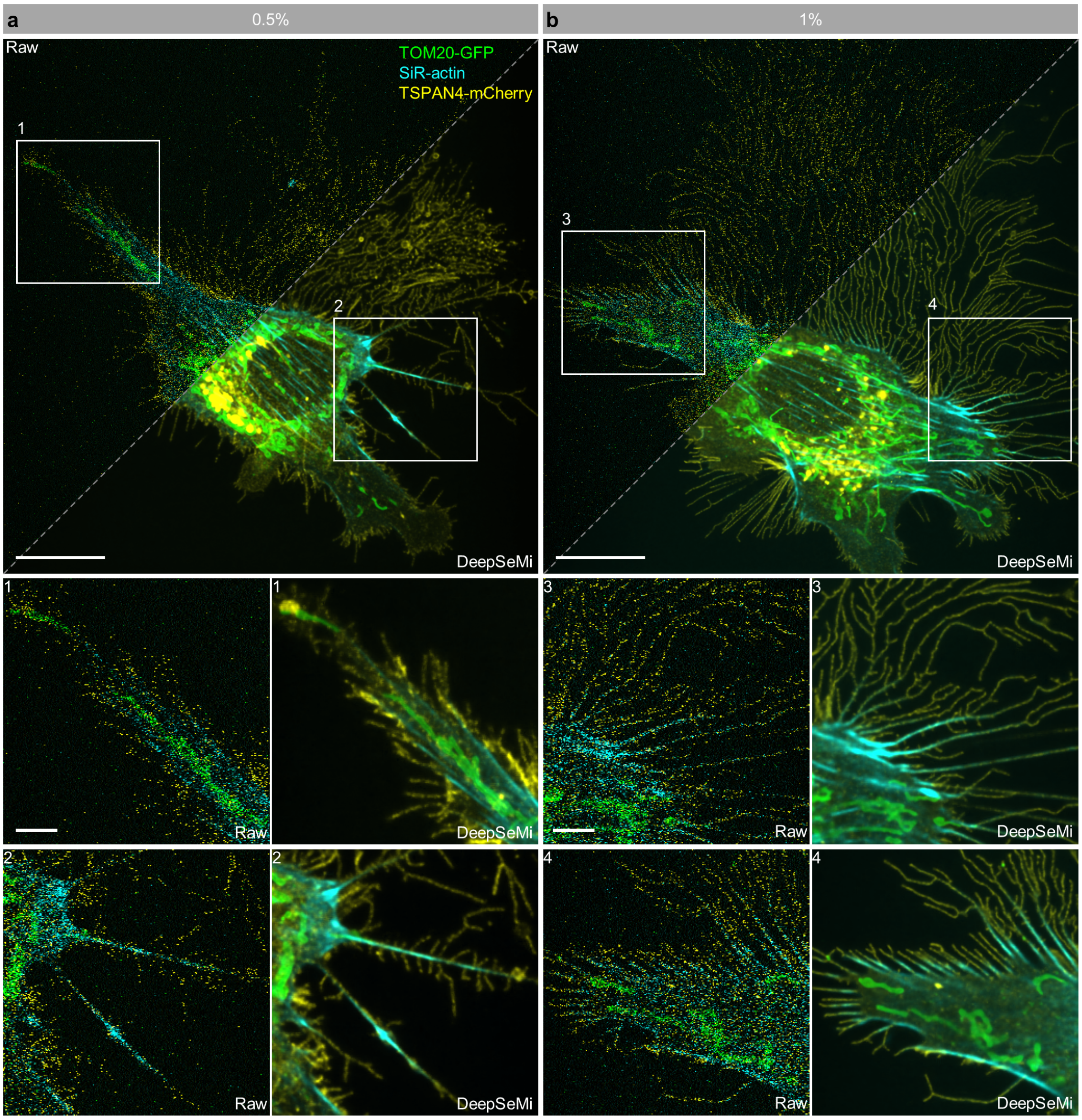


153 **Supplementary Figure 15. DeepSeMi enables high-SNR imaging of tri-color labeled**
154 **L929 cells in low light**. **a-b**, Raw (left) and DeepSeMi-enhanced (right) tri-color labeled
155 L929 cells in 0.5% and 1% laser power (488 nm, 561 nm, and 638 nm), respectively. The
156 second row and the third row presented the zoom-in panels of the white box outlined area
157 in the raw (left) and DeepSeMi-enhanced (right) global view global. Scaler bars are 20
158 µm in the global views, and 5 µm in the zoom-in views.

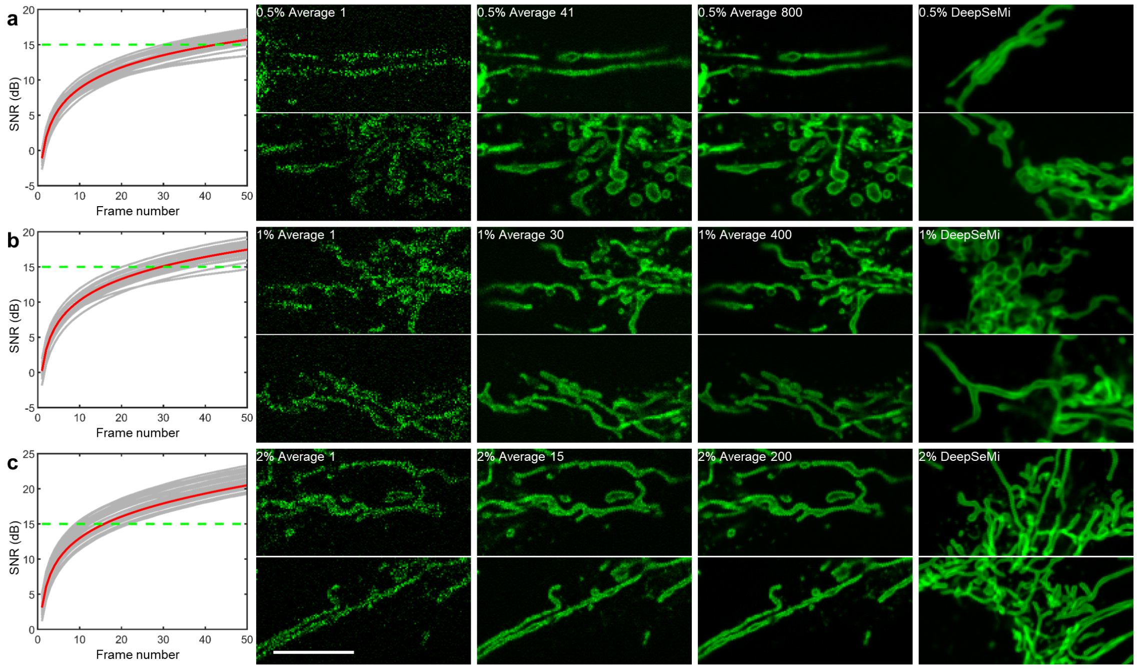


**Supplementary Figure 16. 15-fold increment of photon budgets by DeepSeMi.** We designed a special experiment to calibrate the photon budge enhancement through DeepSeMi in a commercial confocal microscope, which is defined as the multiplication of excitation power in raw captures while reaching the same SNR after DeepSeMi enhancement. To acquire the ground truth image in the experiment, we averaged 800 frames of statistic mitochondria at extremely low excitation laser intensity (0.5% at 488 nm). With the synthetic ground truth, we found that DeepSeMi-enhanced mitochondria with a single frame at 0.5% excitation power reached 15 dB SNR. Towards the bar of 15 dB, we found at least 41 raw frames were required to produce 15 dB SNR through averaging (**a**). Considering the fluorescence yield are linearly proportional to the one-photon excitation power, the number of raw frames used for averaging to some extent represents the multiplication of excitation power for raw capture to catch up with the imaging quality of DeepSeMi enhanced results. We repeated the same procedure for excitation power 1% (**b**) and 2% (**c**). We found the number of enlisted raw frames for reaching 15 dB gradually decayed to 30 frames (**b**) and 15 frames (**c**) as the excitation power increased to 1% and 2%, respectively. Given the fact that DeepSeMi achieves higher SNR as the excitation power, it is safe to state that DeepSeMi increase the photon budget at least 15 times.

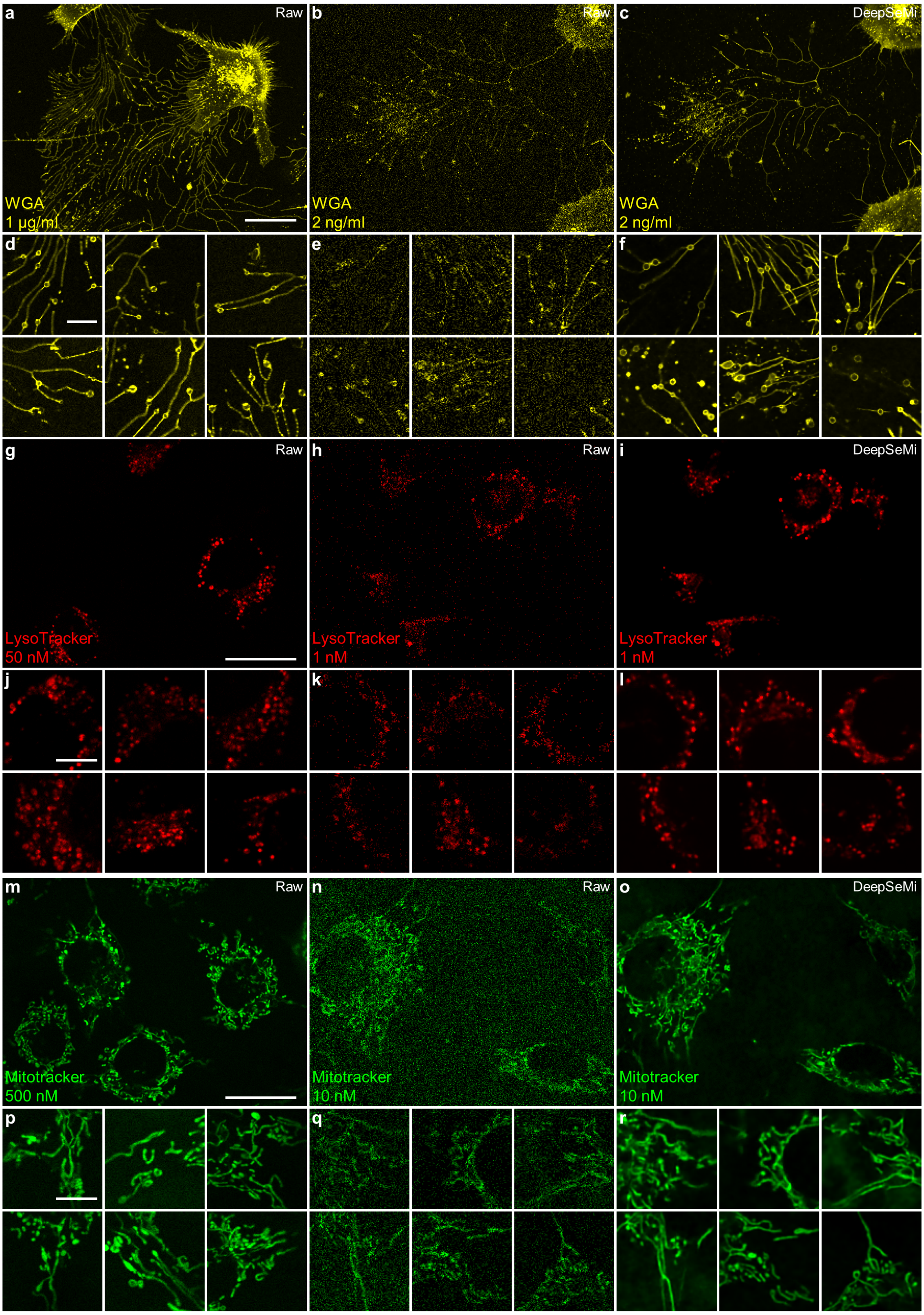


178 **Supplementary Figure 17. DeepSeMi significantly enhances organelle imaging**
179 **results with the dye dilution. a**, The raw confocal imaging results of cells labeled by

WGA with a standard concentration (1 μg/ml). The Nikon A1 was at galvano mode and the excitation laser was at 488 nm (2% power intensity). Scale bar, 30 μm (3 independent trials, each covering more than 10 cells). **b**, Imaging results of the same kind of cells and in the same condition as **a** but labeled by WGA with a diluted concentration (2 ng/ml, 500 times diluted compared to **a**). **c**, DeepSeMi enhanced results of **b**. **d**, Magnified views about migrasomes from raw recordings with standard WGA concentration as in **a**. Scale bar, 10 μm. **e**, Magnified views about migrasomes from raw recordings with 500 times diluted WGA as in **b**. **f**, DeepSeMi enhanced results in **e**. **g**, The raw confocal imaging results of cells labeled by LysoTracker with a standard concentration (50 nM). The excitation laser was at 638 nm (3% power intensity). Scale bar, 30 μm (3 independent trials, each covering more than 10 cells). **h**, The imaging results of the same kind of cells and in the same condition as **g** but labeled by LysoTracker with a diluted concentration (1 ng/ml, 50 times diluted). **i**, The DeepSeMi enhanced results of **h**. **j**, Magnified views about lysosomes from raw recordings with standard LysoTracker concentration in as **g**. Scale bar, 10 μm. **k**, Magnified views about lysosomes from raw recordings with 50 times diluted LysoTracker as in **h**. **l**, DeepSeMi enhanced results in **h**. **m**, The raw confocal imaging results of cells labeled by MitoTracker with a standard concentration (500 nM). The excitation laser was at 561 nm (0.5% power intensity). Scale bar, 30 μm (3 independent trials, each covering more than 10 cells). **n**, The imaging results of the same kind of cells and in the same condition as **m** but labeled by MitoTracker with a diluted concentration (10 ng/ml, 50 times diluted). **o**, The DeepSeMi enhanced results of **n**. **p**, Magnified views about mitochondrial from raw recordings with standard MitoTracker concentration as in **m**. Scale bar, 10 μm. **q**, Magnified views about mitochondrial from raw recordings with 50 times diluted MitoTracker concentration as in **n**. **r**, DeepSeMi enhanced results in **n**.

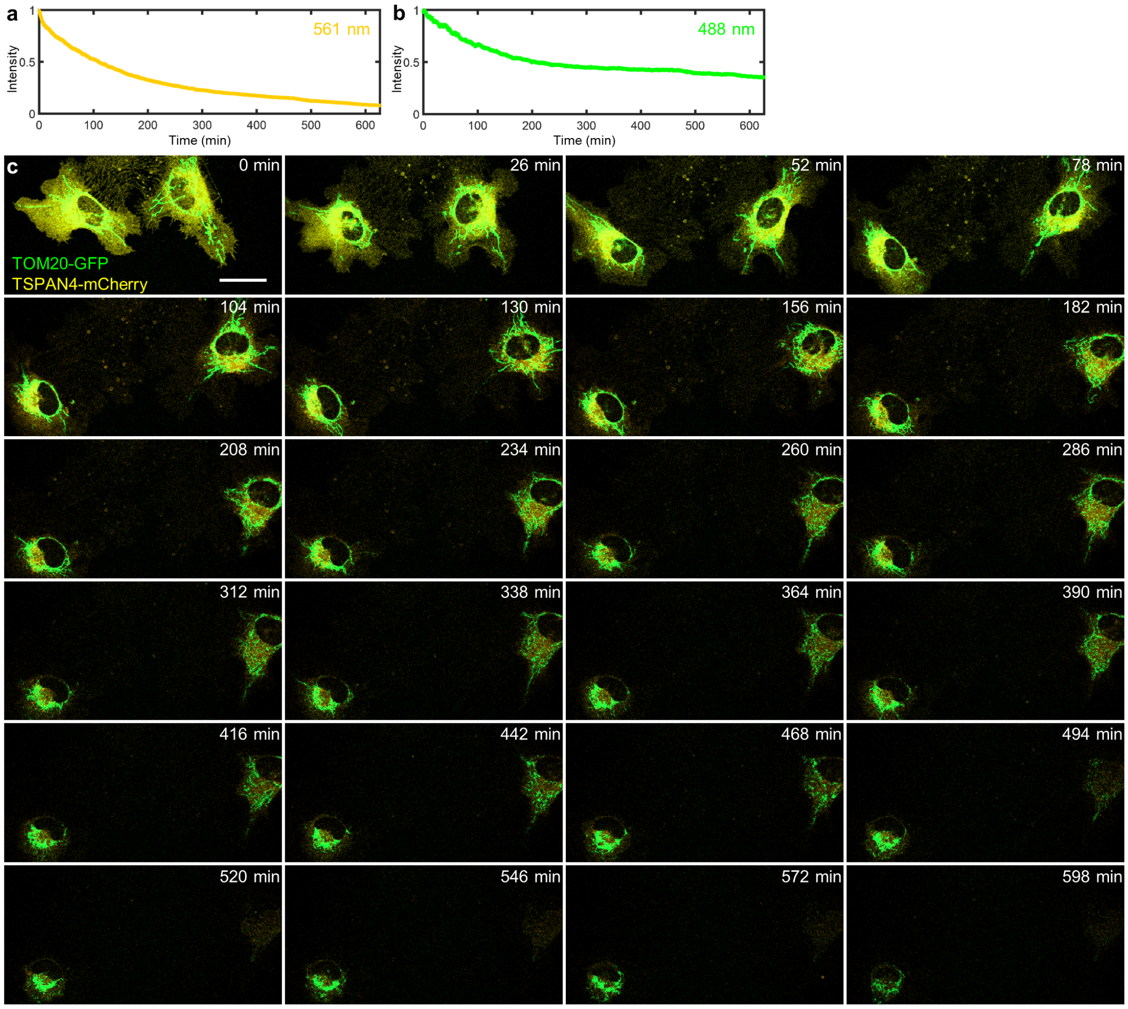


**Supplementary Figure 18. Significant photobleaching induced by dual-color confocal imaging. a-b**, Normalized fluorescence intensity over continuous 10 hours confocal imaging under 488 nm (right) and 561 nm excitation (left), respectively. 0.36 mW 488 nm and 0.44 mW 561 nm lasers were used for imaging dual-labeled L929 cells at 1.92Hz. **c**, Exemplary time-lapse dual-color images at different time points over 10 hours of imaging clearly show photobleaching. All images are raw captured without DeepSeMi enhancement. Scale bar, 30 um.

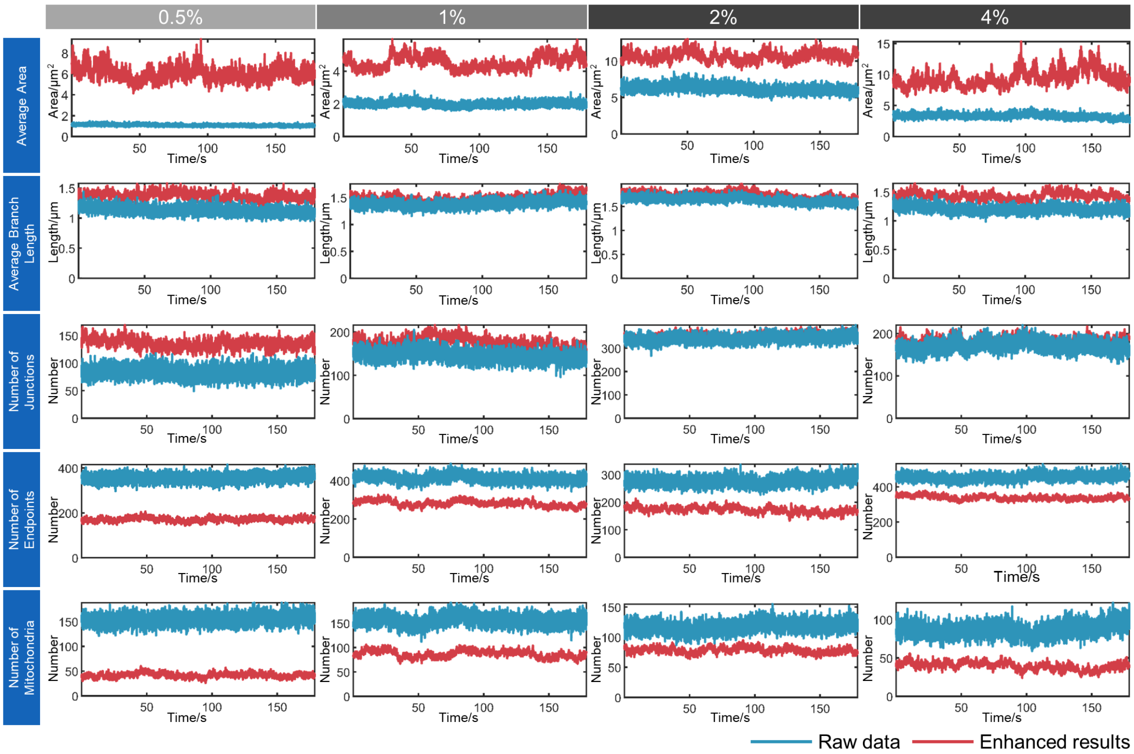


**Supplementary Figure 19. Statistics of mitochondrial segmentation and skeletonization under different illumination powers with and without DeepSeMi enhancement.** The average area, branch length, number of junctions, number of endpoints, and number of mitochondria were calculated before (blue) and after DeepSeMi enhancement (red) for a 180-seconds imaging session at 30 Hz. Since the DeepSeMi effectively reunited fragment mitochondria under noise contamination, the average area, branch length, and the number of junction points are increased after DeepSeMi enhancement, while the number of endpoints and the number of mitochondria are accordingly reduced.

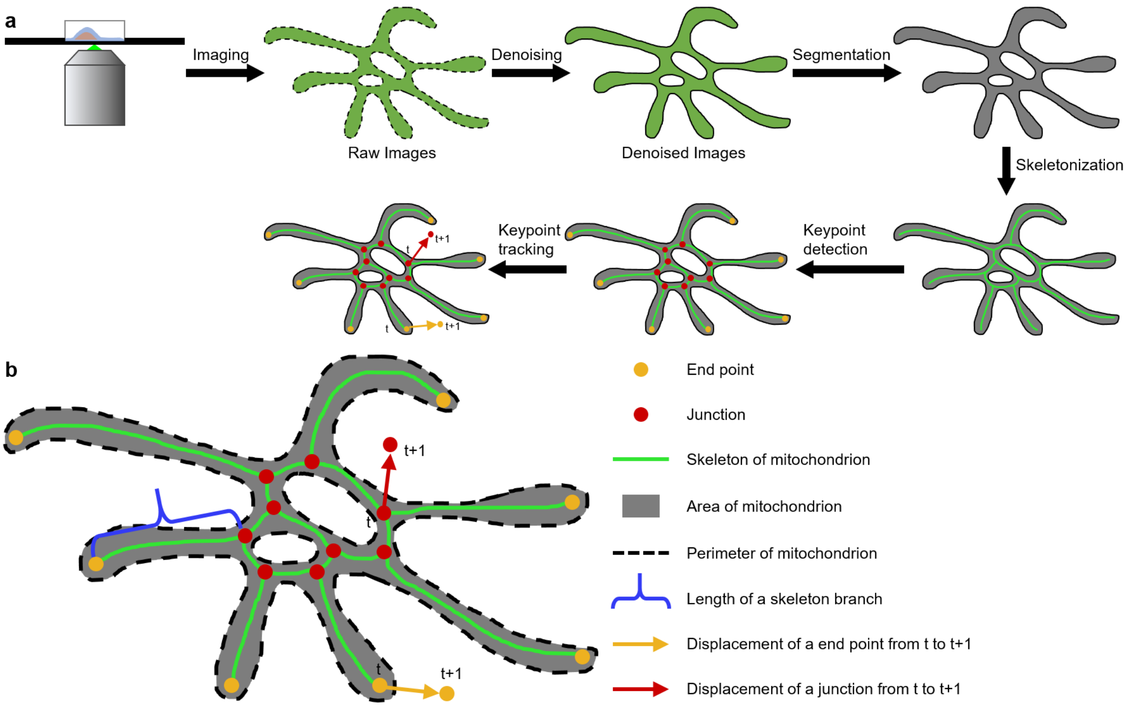


**Supplementary Figure 20. Automated analysis of recorded mitochondria with DeepSeMi enhancement**. **a**, The captured videos of mitochondria are firstly denoised by DeepSeMi to remove noise contaminations, and then segmented with a simulation-supervision machine learning algorithm [7]. The binary mask is then skeletonized (Methods) with key points (including end point, junction point) highlighted. Those key points are used for tracking the motion of the mitochondria. **b**, Illustrations of extracted features and associated measurements from the mitochondria, including end point, junction point, skeleton, area, perimeter, and branch length.

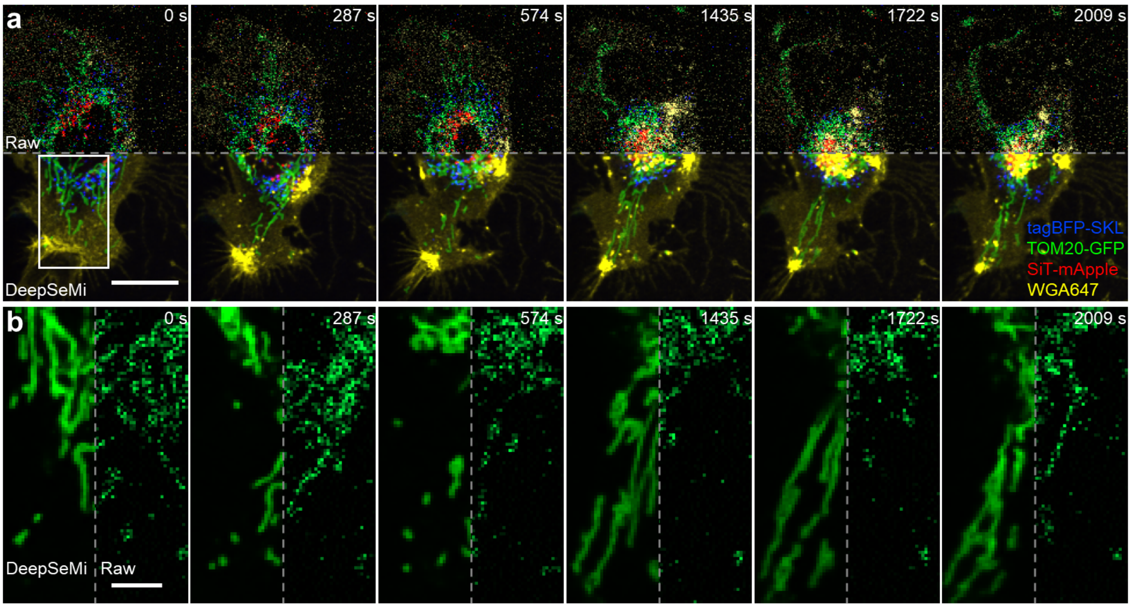


**Supplementary Figure 21. DeepSeMi-enhanced imaging results of L929 cells treated with Lat-A. a**, Raw (top) and DeepSeMi-enhanced (bottom) long-term imaging of L929 cells with four organelles labeled colorfully (TOM20–GFP, WGA647, TagBFP–SKL, and SiT-mApple). Latrunculin-A (lat-A) was added to the cell culture medium at 0s to decompose the cytoskeleton (Methods). Scale bar 20 μm. **b**, Mitochondria deformation during 33 minutes-long time-lapse imaging after treatment with lat-A. For each panel, the left part represents DeepSeMi enhancement and the right panel represents the raw image. Scale bar 5 μm.

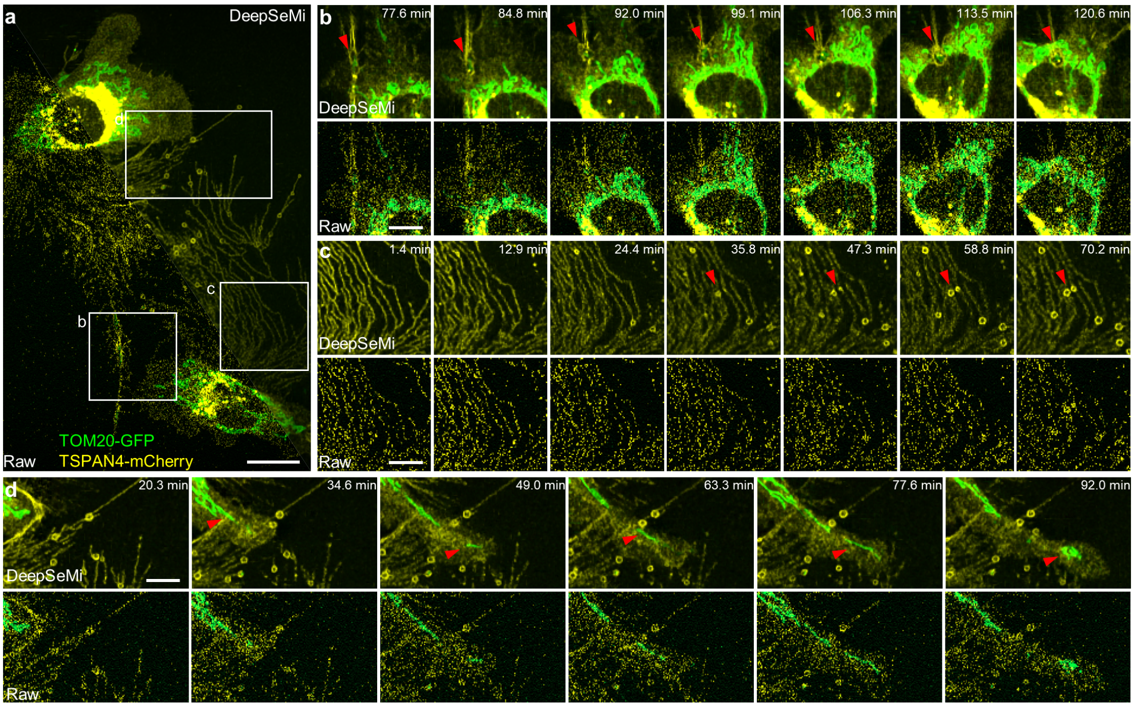


**Supplementary Figure 22. DeepSeMi unveiled migrating cells interacting with a migrasome, producing migrasomes, and expelling mitochondria in a low light dosage. a**, Raw (left) and DeepSeMi enhanced (right) observation of two-cell interactions. Cells with mitochondria (green, TOM20-GFP) and migrasomes (yellow, TSPAN4-mCherry) labeled were imaged at 95.1 μW for 2 hours at 1.16 Hz. Scale bar, 20 μm. **b**, Time-lapse process of cell interacting with a migrasome (marked by red arrows) during migration by DeepSeMi enhanced (top) and raw (bottom) captures. The migrasome is almost invisible in the raw movie. Scale bar, 10 μm. **c,** Time-lapse process of a cell producing migrasomes during migration by DeepSeMi enhanced (top) and raw (bottom) captures. The retraction fibers were produced when the cell was crawling, and spherical migrasomes (marked by red arrows) were generated on the retraction fibers by the regulation of the cell. Scale bar, 10 μm. **d**, Time-lapse process of cell expelling mitochondria (marked by red arrows) during migration by DeepSeMi enhanced (top) and raw (bottom) captures. Scale bar, 10 μm.

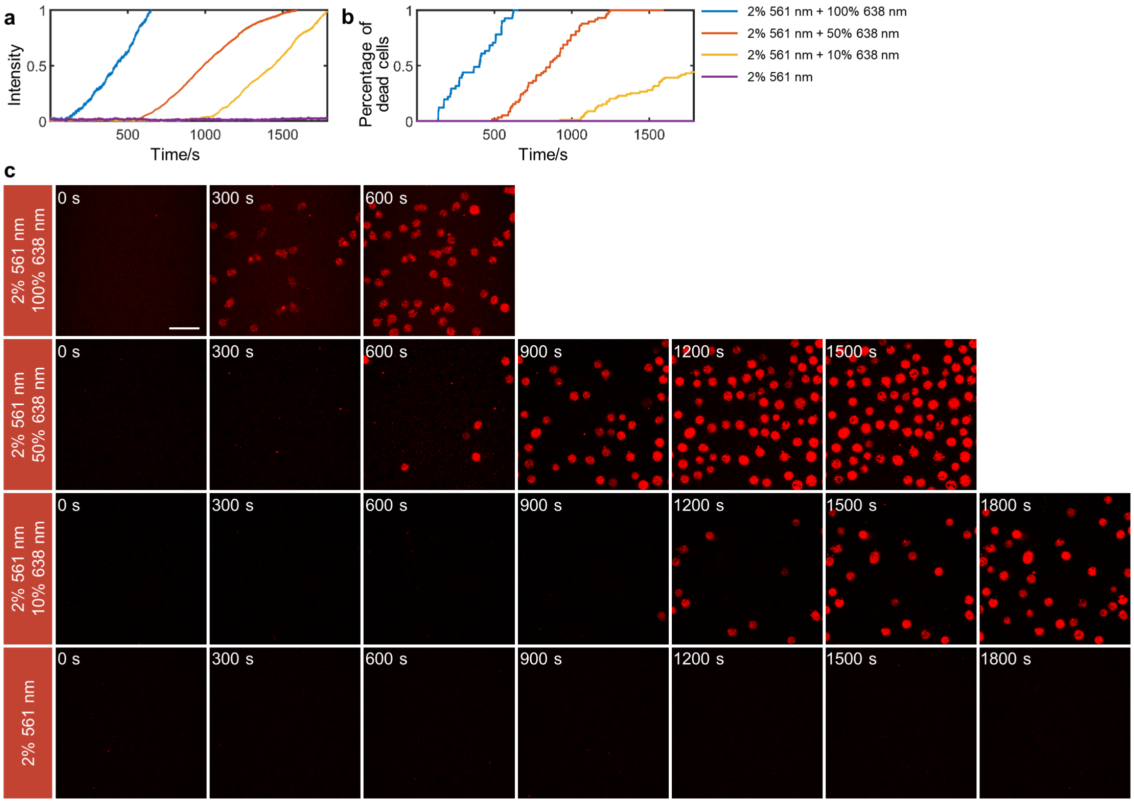


**Supplementary Figure 23. Evaluation of phototoxicity in imaging *Dictyostelium* cells.** The *Dictyostelium* cells were simultaneously illuminated by the 638 nm laser and the 561 nm laser. The 638 nm laser was used for generating phototoxicity on *Dictyostelium* cells. The 561 nm laser was used for imaging to evaluate the phototoxicity brought by 638 nm illumination. When *Dictyostelium* cells died because of phototoxicity, the permeability of the membrane changed and the propidium iodide in the micro-environment entered into *Dictyostelium* cells which facilitated fluorescence imaging under 561 nm excitation. Four laser powers at 640 nm were assessed (0%, 10%, 50%, 100%). **a**, Statistics of fluorescence intensity change under 561 nm excitation during imaging as a function of time in four conditions. The fluorescence intensity is normalized into 0 to 1. **b**, The number of *Dictyostelium* cell deaths during imaging as a function of time in four conditions. The number of cell deaths is normalized into 0 to 1, which is highly correlated with curves in **a**. **c**, Exemplary results of fluorescence imaging of *Dictyostelium* cells under 561 nm excitation during phototoxicity experiments at different time points. Scale bar, 20 μm.

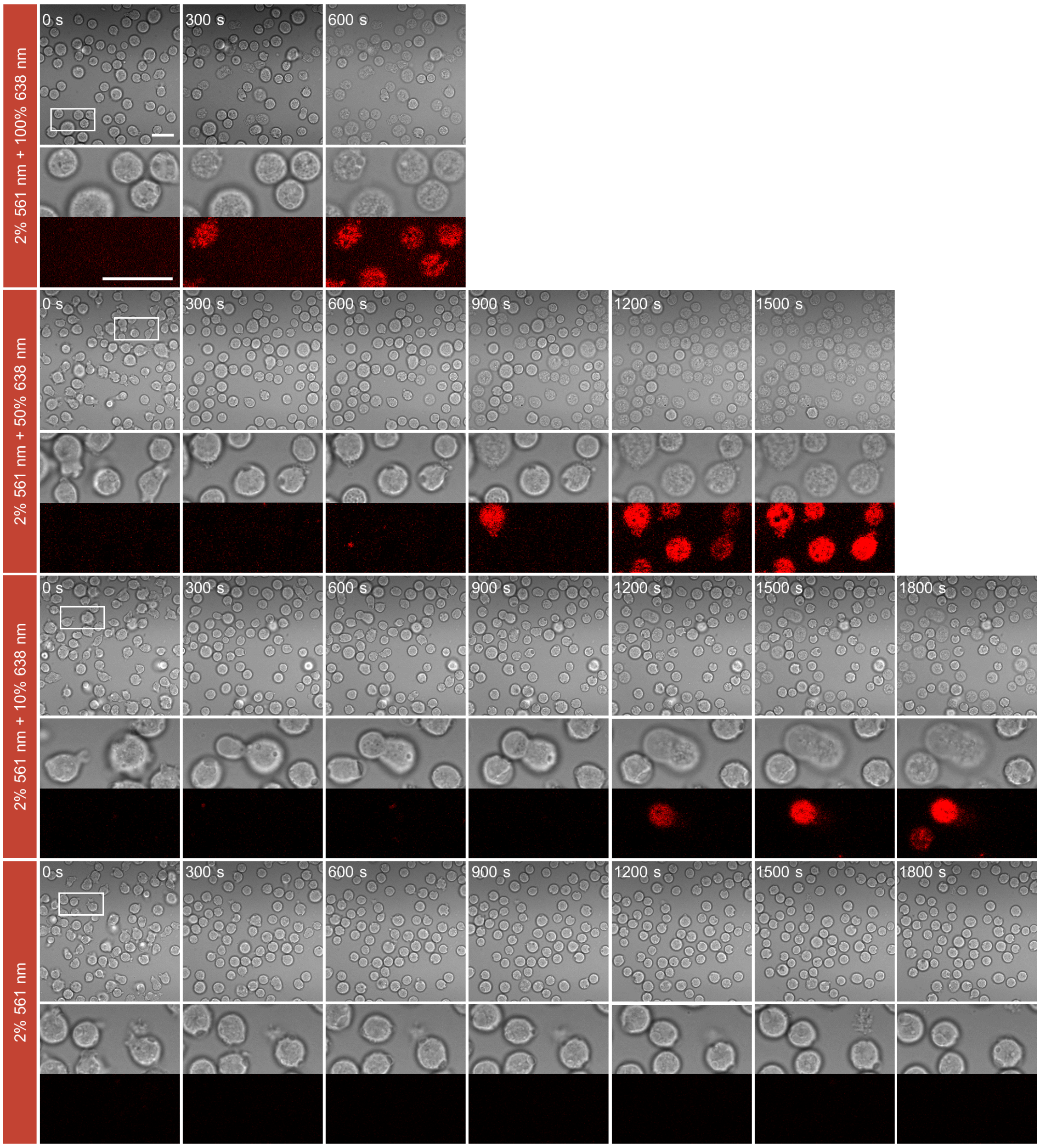


**Supplementary Figure 24. Evaluation of phototoxicity in imaging *Dictyostelium* cells with a bright-field microscope imaging.** The imaging properties are the same as Supplementary Figure 23. The boundaries of the cells lose sharpness as the cell suffers phototoxicity, which provides another clue to monitor the health status of the cell. Four laser powers at 638 nm were assessed. For each power, the first row shows a global view of the bright field microscope capture, and the top part of the second row shows a zoom-in image of the white box in the global view, and the bottom part of the second row shows the corresponding fluorescence image under 561 nm excitation. Scale bar, 20 µm.

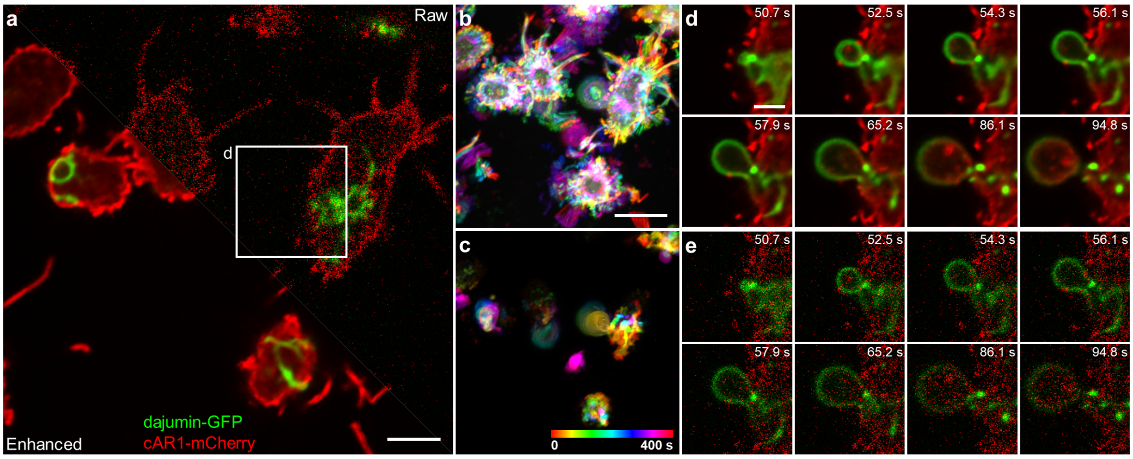


**Supplementary Figure 25. DeepSeMi enables high-SNR imaging of contractile vacuole dynamics in photosensitive *Dictyostelium* cells**. **a**, DeepSeMi enhanced (left) and raw (right) images of *Dictyostelium* cells, where membranes are labeled in red and contractile vacuoles are labeled in green. 6000 frames are recorded in 400 seconds. Scale bar, 5 μm. **b-c**, Temporal-color coded DeepSeMi enhanced and raw images in **a**, respectively. Scale bar, 10 μm. **d-e**, Recorded time-lapse process of a contractile vacuole generation enclosed by the white box in **a** by DeepSeMi enhanced and raw images, respectively. Scale bar, 3 μm.

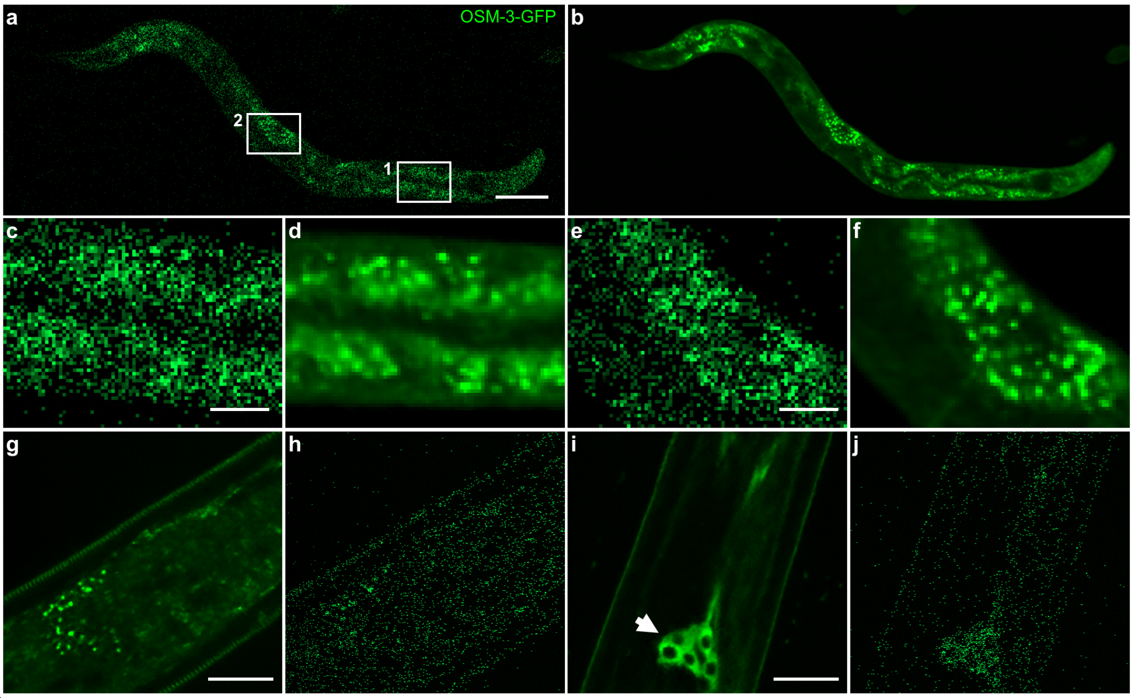


**Supplementary Figure 26. DeepSeMi enhanced cellular observation in scattering *C. elegans* in vivo. a-b**, *In vivo* imaging of *C. elegans* in a millimeter-scale field-of-view (FOV) with a 10× objective by raw and DeepSeMi-enhanced captures, respectively. Scale bar, 100 µm. **c-d**, Zoom-in panels in the white box ("1") outlined area in **a**, respectively. Scale bar 20 µm. **e-f**, the same as **c-d** but for the white box ("2"). Scale bar 20 µm. **g-h**, *C. elegans* imaging with a 100× objective by DeepSeMi enhanced and raw captures, respectively. Scale bar, 15 µm. **i-j**, Position where the hole shape structure was clearly recovered through DeepSeMi. Scale bar, 15 µm.

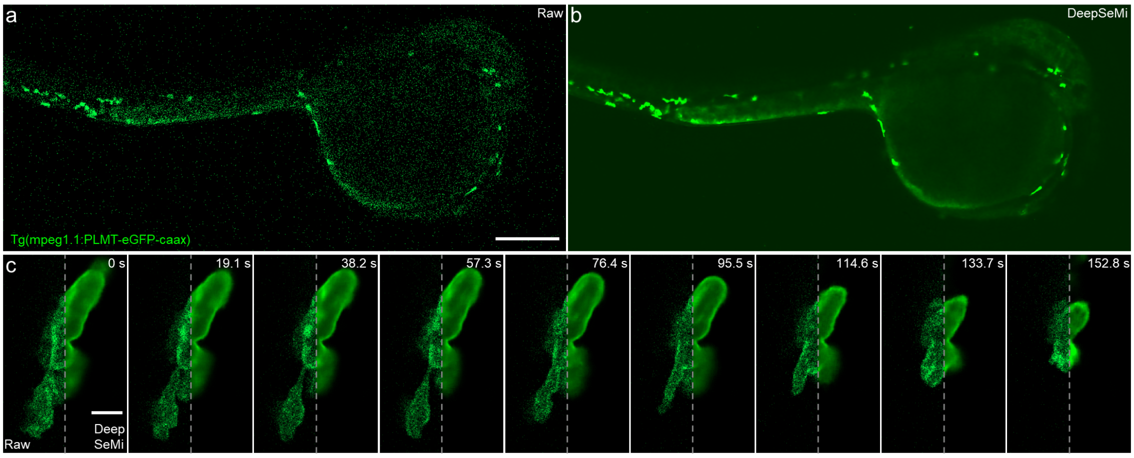


**Supplementary Figure 27. DeepSeMi enhances observation of zebrafish larvae in a low light dosage**. **a-b**, Raw and DeepSeMi enhanced observation of zebrafish larvae, respectively. The larva was observed in a commercial confocal microscope with a low magnification objective (10×, NA 0.45). Scale bar, 200μm. **c**, Time-lapse cellular imaging of macrophage in the zebrafish through a high magnification objective (100×, NA 1.45). For each time point, the left part is raw image and the right part is DeepSeMi enhanced image. Scale bar, 5μm.

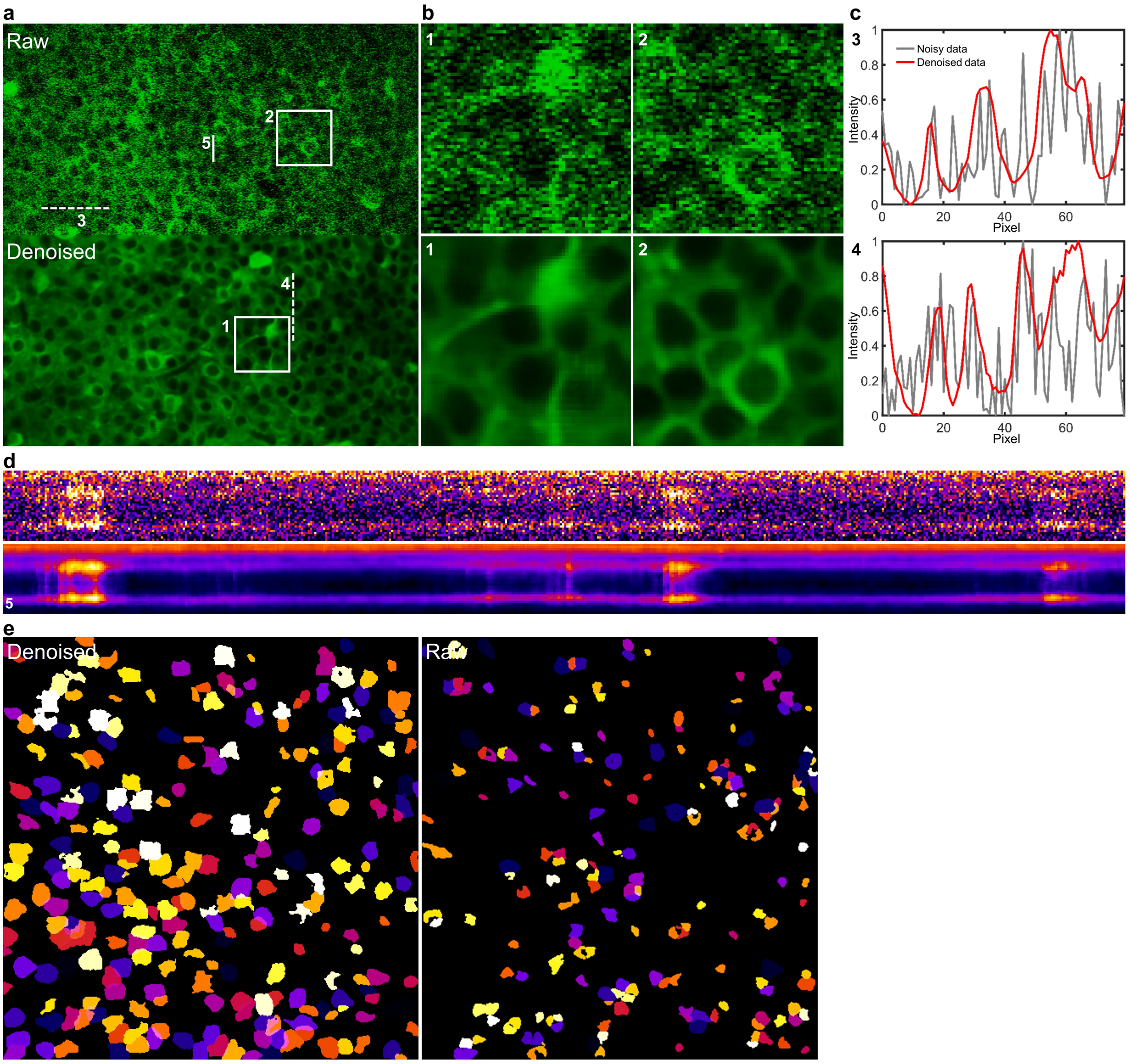


**Supplementary Figure 28. DeepSeMi effectively recovers functional data on open-sourced two-photon Neurofinder datasets**. **a**, Comparison of denoising results of DeepSeMi (bottom) with raw frame (top) in Neurofinder datasets [9]. The neuronal structures are clearly recovered by DeepSeMi. **b**, Zoom-in panels of the white box outlined regions in a. **c**, Cross-sectional intensity profiles along the white dashed lines in **a**, where red represents the DeepSeMi denoising and the gray represents raw data. **d**, Kymographs (*x-t* data) of raw data (top) and DeepSeMi denoised data (bottom) origins from the white solid line in **a**. **e**, Neuron segmentation results of raw (right) and DeepSeMi denoised data (left) through open-source CaImAn package [9]. 196 neurons were extracted from the raw image, and 252 neurons were extracted from the DeepSeMi denoised image.

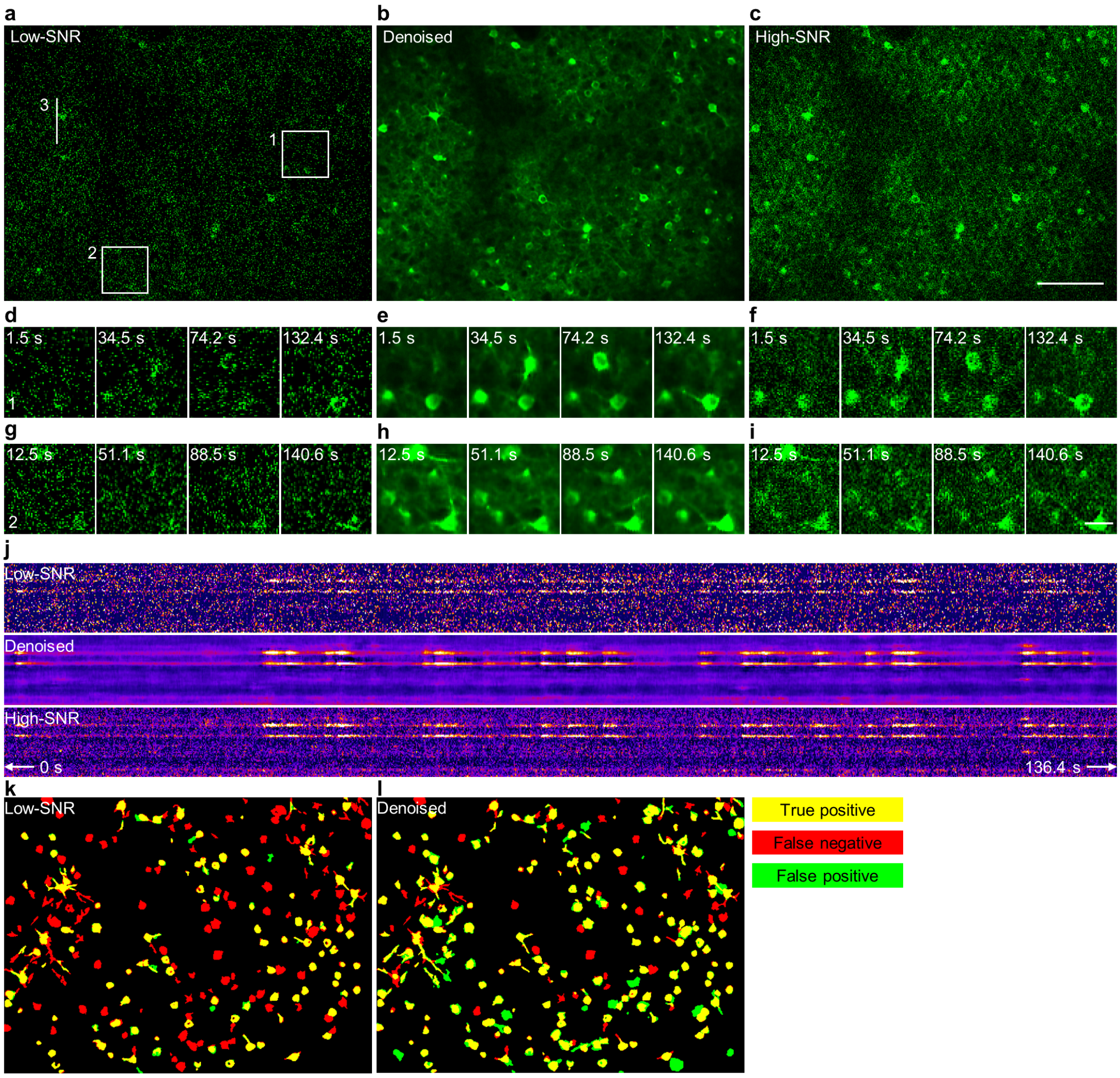


**Supplementary Figure 29. Evaluation of DeepSeMi on hybrid high and low-SNR functional imaging.** We set up a hybrid two-photon microscope with two-channels where one achieves 10-fold SNR compared to the other one. **a-c**, Left to right, low SNR image, DeepSeMi restored image, the high SNR image as a reference. Scale bar, 100 μm. **d-i**, Zoom-in panels of the white box outlined regions in **a-c** at different time points. Scale bar, 20 μm. **j**, Kymographs (x-t data) from the lines in **a-c**. **k-i**, The CNMF segmentation results of low SNR images, restored images and high SNR images. True positives, false positives, and false negatives are annotated.

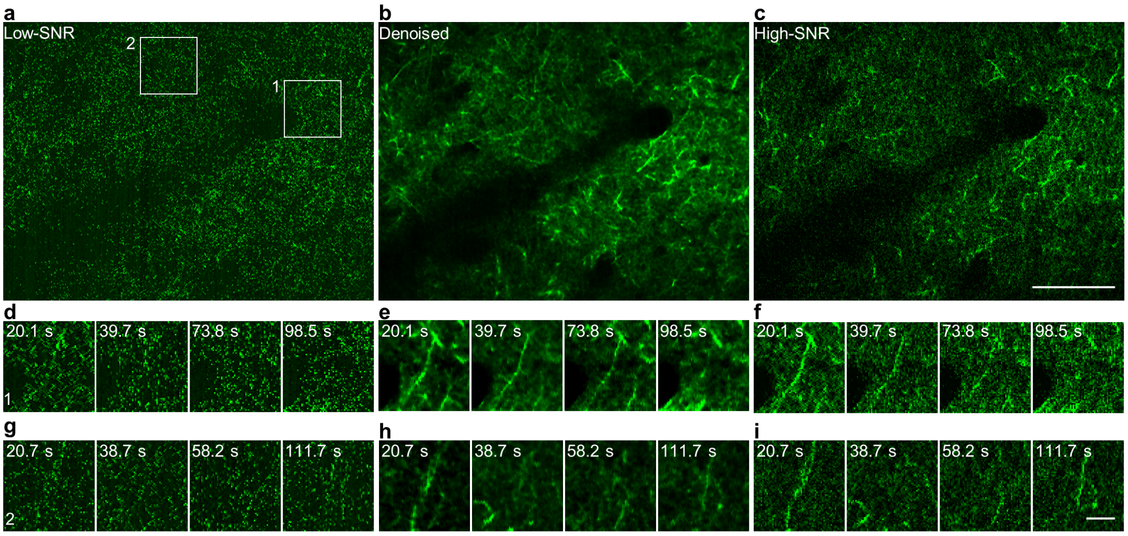


**Supplementary Figure 30. Evaluation of DeepSeMi on hybrid high and low-SNR dendritic imaging**. **a-c**, Left to right, low SNR image, DeepSeMi restored image, the high SNR image as a reference. Scale bar, 100 µm. **d-i**, Zoom-in panels of the white box outlined regions in **a-c** at different time points. Scale bar, 20 µm.
